# Disputing space-based biases in unilateral complex regional pain syndrome

**DOI:** 10.1101/2020.01.02.893263

**Authors:** Monika Halicka, Axel D Vittersø, Hayley McCullough, Andreas Goebel, Leila Heelas, Michael J Proulx, Janet H Bultitude

**Author notes:** Corresponding author: Monika Halicka, Department of Psychology, University of Bath, Claverton Down Road, Bath BA2 7AY, United Kingdom. **Abbreviations** BPDS = Bath CRPS Body Perception Disturbance Scale; BPI = Brief Pain Inventory; CRPS = Complex Regional Pain Syndrome; EHI = Edinburgh Handedness Inventory; MNLB = Mental Number Line Bisection; POMS = Profile of Mood States; PSE = Point of Subjective Equality; PSS = Point of Subjective Simultaneity; TOJ = Temporal Order Judgement; VF = Visual Field.

## Abstract

There is some evidence that people with Complex Regional Pain Syndrome (CRPS) show reduced attention to the affected relative to unaffected limb and its surrounding space, resembling hemispatial neglect after brain injury. These neuropsychological symptoms could be related to central mechanisms of pathological pain and contribute to its clinical manifestation. However, the existing evidence of changes in spatial cognition is limited and often inconsistent. We examined visuospatial attention, the mental representation of space, and spatially-defined motor function in 54 people with unilateral upper-limb CRPS and 22 pain-free controls. Contrary to our hypotheses and previous evidence, individuals with CRPS did not show any systematic spatial biases in visuospatial attention to or representation of the side of space corresponding to their affected limb (relative to the unaffected side). We found very little evidence of directional slowing of movements towards the affected relative to unaffected side that would be consistent with motor neglect. People with CRPS were, however, slower than controls to initiate and execute movements with both their affected and unaffected hands, which suggests disrupted central motor networks. Finally, we found no evidence of any clinical relevance of changes in spatial cognition because there were no relationships between the magnitude of spatial biases and the severity of pain or other CRPS symptoms. The results did reveal potential relationships between CRPS pain and symptom severity, subjective body perception disturbance, and extent of motor impairment, which would support treatments focused on normalizing body representation and improving motor function. Our findings suggest that previously reported spatial biases in CRPS might have been overstated.

## 1. Introduction

Growing evidence supports the notion that chronic pain is a disease of the central nervous system involving functional and structural reorganisation of the brain (for reviews, see Henry, Chiodo, & Yang, 2011; Lee, Nassikas, & Clauw, 2011; Seifert & Maihöfner, 2008). One condition in which such reorganisation has been observed is Complex Regional Pain Syndrome (CRPS), a disorder that can affect one or more limb(s) and involves pain and other sensory, motor, and autonomic symptoms that are disproportionate to any inciting injury. Abnormal higher-order cortical processing in CRPS is further evidenced by cognitive changes in the representation of and attention to the CRPS-affected limb and the corresponding side of external space (e.g. Bultitude, Walker, & Spence, 2017; Legrain, Bultitude, De Paepe, & Rossetti, 2012; Lewis, Kersten, McCabe, McPherson, & Blake, 2007; Moseley, Gallace, & Spence, 2009; Schwoebel, Coslett, Bradt, Friedman, & Dileo, 2002). These changes have been referred to as “neglect-like”, because they resemble those typical of hemispatial neglect that can occur after brain injury. For example, deficits in attention to or representation of the affected (relative to unaffected) side have been found on tests of tactile attention (Moseley et al., 2009, 2012; Reid et al., 2016), visual attention (Bultitude et al., 2017; Filbrich et al., 2017), and the mental representation of space (Sumitani et al., 2014). Furthermore, similar to people with hemispatial neglect after brain injury, people with CRPS reported or presented with underutilisation of the affected limb during spontaneous movements that could not be fully explained by primary motor deficits (Galer et al., 1995; Galer & Jensen, 1999). Systematic measurement of motor performance of people with CRPS revealed slower and more variable movements when they used their affected hand, but also when movements were performed in the affected side of space regardless of which hand they used (Reid et al., 2018). Thus, there is evidence that people with CRPS can show space-based neuropsychological changes that resemble perceptual, representational, and motor neglect.

Previous studies suggest that there is a relationship between spatial biases and the manifestation and maintenance of CRPS symptoms. For example, the severity of self-reported “neglect-like” symptoms (Frettlöh et al., 2006) was associated with greater pain intensity, worse long-term pain outcomes, sensory loss, and motor impairment in the affected limb (Frettlöh et al., 2006; Kolb et al., 2012; Wittayer et al., 2018). Also, the magnitude of perceptual and motor spatial biases on experimental tasks correlated with greater pain intensity and longer CRPS duration (Reid et al., 2016, 2018). Furthermore, larger temperature asymmetry between the affected and unaffected arms was related to greater magnitude of tactile spatial attention bias (Moseley et al., 2009, 2012). This asymmetry was reduced by resting the affected hand in the unaffected side of space, and by using prismatic lenses to produce the illusion of such positioning (Moseley et al., 2012, 2013). Reduction of pain and other CRPS symptoms following prism adaptation treatment (Bultitude & Rafal, 2010; Christophe, Chabanat, et al., 2016; Sumitani, Rossetti, et al., 2007), which is thought to increase attention to the affected side relative to the unaffected side, further supports the clinical relevance of spatial attention in CPRS. Therefore, understanding spatial biases in CRPS, and how they relate to clinical symptoms, could provide insights into the prevention and treatment of the disorder.

Some findings, however, have called into question the presence of spatial biases in CRPS, or the extent to which they resemble hemispatial neglect. There is a share of research showing no evidence of spatial biases in CRPS (e.g. Christophe, Chabanat, et al., 2016; Filbrich et al., 2017; Filippopulos, Grafenstein, Straube, & Eggert, 2015; Förderreuther, Sailer, & Straube, 2004; Kolb et al., 2012; Reid et al., 2016; Reinersmann et al., 2012; Wittayer et al., 2018). Furthermore, some researchers found that people with CRPS presented with spatial biases in the *opposite* direction to what would be considered “neglect-like”. The best example of such findings is that the representation of external space relative to one’s body was shifted towards the CRPS-affected side in several group studies (Sumitani et al., 2014; Sumitani, Rossetti, et al., 2007; Sumitani, Shibata, et al., 2007; Uematsu et al., 2009). Two case reports also described a CRPS patient who consistently showed higher attention to her affected side relative to her unaffected side across a battery of tests of spatial cognition (Christophe, Delporte, et al., 2016; Jacquin-Courtois et al., 2017). Some of these negative findings could be due to insufficient sensitivity of the tests used. For example, subtle spatial biases in people with CRPS, who typically do not have any brain injury, might not be evident on tasks such as classic pen-and-paper line bisection. Small sample sizes, in combination with the known heterogeneity of CRPS presentation (Bruehl et al., 2016; Marinus et al., 2011), could also account for some of the inconsistencies between studies regarding the observed presence or direction of spatial biases. Furthermore, very few previous studies tested different aspects of neglect (i.e. perceptual, representational, and motor) in the same group of participants (see Christophe, Chabanat, et al., 2016; Reid et al., 2016; Sumitani et al., 2014 for exceptions). Therefore, it can be difficult to ascertain if discrepancies between the demonstrated spatial biases (or lack thereof) across studies are due to differences between the participants, or because CRPS can affect one aspect of spatial cognition and not another. The aim of the present study was therefore to use sensitive measures to examine multiple aspects of spatial cognition in a large sample of individuals with CRPS.

We designed a battery of tests based on established approaches used to measure hemispatial neglect following a stroke, as well as pseudoneglect (the mild leftward bias that is commonly found in groups of healthy participants when performing certain spatial tasks; Jewell & McCourt, 2000). We used three experimental tasks to test for visuospatial biases. The first, the Temporal Order Judgement (TOJ) task, requires participants to indicate the relative timings of pairs of spatial stimuli (one presented in each visual field). The TOJ measures covert spatial attention, based on the premise that information that is subject to greater attention is perceived earlier relative to information that is subject to lesser attention (Spence & Parise, 2010). The second, the Landmark task, requires participants to judge the relative distance of two stimuli and is thought to measure visuospatial representations (Makin et al., 2010). A tendency to underestimate the distance on one side of space (relative to the other) would be consistent with diminished visual representation of that side. The third, the Greyscales task (Nicholls et al., 1999), requires participants to judge the relative luminance of two equally shaded greyscale stimuli that are arranged one above the other such that one stimulus has greater luminance on the left and one has greater luminance on the right. Participants with an attention bias will tend to show greater reliance on the luminance difference on one side of the stimulus display when making their decision. This tests for any bias in overt spatial attention without posing temporal demands on the task. In addition to the tests of visuospatial biases, our battery also included a Mental Number Line Bisection (MNLB) task designed to measure any biases in mental representations of space. This is based on the existing evidence that numbers are mentally represented in a linear arrangement (with smaller numbers located to the left, and larger numbers to the right side of space; Dehaene, Bossini, & Giraux, 1993). Thus, individuals with biased mental representation of space will tend to underestimate or overestimate the midpoint of number intervals on a mental number line. Finally, to test for any spatial biases in motor function, we measured the speed of movement initiation and execution when participants reached from different starting locations towards targets appearing either in the affected or unaffected side of space. Slower initiation of movements directed towards the affected relative to unaffected side of space, even with the unaffected hand, defines directional hypokinesia towards the affected side, whereas slower execution of the same movements defines directional bradykinesia. This test of spatially-defined motor function was identical to that used previously to measure directional hypokinesia in right-hemisphere stroke patients (Sapir et al., 2007). We hypothesised that participants with CRPS (compared to pain-free controls) would present with spatial biases in attention to and representations of the affected side of space, and with slowed initiation and execution of movements directed towards the affected relative to unaffected side of space.

In addition to evaluating group differences in spatial cognition, we also explored relationships between pain, CRPS severity, and the extent of any neuropsychological changes in people with CRPS. Previously discussed literature shows that changes in spatial cognition correlate with clinical features of CRPS such as pain intensity, sensory and motor impairment, and temperature asymmetry (Frettlöh et al., 2006; Kolb et al., 2012; Moseley et al., 2009, 2012, 2013; Reid et al., 2016, 2018; Wittayer et al., 2018). There is also evidence to suggest that they are associated with other cognitive abnormalities (e.g. body perception disturbance; Bultitude et al., 2017) and psychological distress (e.g. depression, anxiety; Michal et al., 2016; Wittayer et al., 2018). However, the relationships between spatial biases and clinical CRPS symptoms are not consistently found (Bultitude et al., 2017; Filbrich et al., 2017; Frettlöh et al., 2006; Michal et al., 2016; Reid et al., 2016; Reinersmann et al., 2012; Vittersø et al., 2019). Considering that some potentially relevant outcomes might be overlooked in the existing literature, we explored our data for relationships between spatial biases and a broad range of participant characteristics (such as age, CRPS duration, and change in hand preference), clinical outcomes (sensory, motor, and autonomic function), self-reported pain, and psychological factors (body perception disturbance, pain-related fear of movement, and mood disturbance). We hypothesised that the magnitude of any observed neuropsychological symptoms would be related to the severity of clinical signs of CRPS.

## 2. Methods

We report how we determined our sample size, all data exclusions (if any), all inclusion/exclusion criteria, whether inclusion/exclusion criterial were established prior to data analysis, all manipulations, and all measures in the study. This study involved a single study visit that was a part of a randomised controlled trial to evaluate the effects of prism adaptation on pain and severity of CRPS symptoms (CRPS PRISMA Trial, ISRCTN46828292, prospectively registered at https://doi.org/10.1186/ISRCTN46828292; see Halicka, Vittersø, Proulx, & Bultitude, 2019 for the trial protocol and analysis plan; note that analysis plan was not pre-registered prior to the research being conducted, but the manuscript was submitted for publication before commencement of any data analysis). The same participants also completed a hand laterality recognition task, which will be reported elsewhere as this was designed to measure lateralised body representation distortion rather than spatial cognition per se. The data reported in this article were collected prior to any trial-related intervention. During the single study visit, participants completed self-report questionnaires; underwent assessment of sensory, motor, and autonomic function; and completed experimental tests of neuropsychological function. The study visit lasted between two to four hours, including breaks between the assessments. The results of the randomised controlled trial will be reported elsewhere. All procedures were carried out in accordance with the Declaration of Helsinki and received ethical approval from National Health Service Oxfordshire Research Ethics Committee A (ref. 12/sc/0557).

Participants were recruited through the National CRPS-UK Registry, internal registry of the Walton Centre NHS Foundation Trust, Oxford University Hospitals NHS Foundation Trust, and other NHS clinics in the UK, word of mouth, advertisements on the funder’s and research centre’s websites, and social media. Participants were screened for eligibility through a telephone interview. To obtain a sample size meeting the trial requirements (21 patients completing the trial for each of the two treatment groups; Halicka et al., 2019), we enrolled 54 adults with CRPS-I affecting primarily one upper limb for at least three months, who met the Budapest research diagnostic criteria at the time of testing (Harden et al., 2010). The control sample consisted of 22 adults without current or chronic pain, who were matched to 22 individual participants with CRPS (i.e. size of one treatment group) by sex, self-reported handedness, and age (±5 years). One limb of each control participant was labelled as the “matched” (i.e. “affected”) limb according to the affected limb of their matched participant with CRPS. Note that although sample sizes are imbalanced, we would expect more variability in the heterogeneous (but larger) sample of participants with CRPS than in healthy control participants (Rusticus & Lovato, 2014). All participants enrolled in the study had no history of neurological disorders, no severe psychiatric disorders that might be associated with perceptual changes (e.g. schizophrenia), were not legally blind, and had sufficient English language ability to provide informed consent. Written informed consent was obtained from all participants prior to any study-related procedures. Participants completed their study visit at the Universities of Bath (36 participants with CRPS, all controls), Liverpool (10 participants with CRPS), or at participants’ homes if unable to travel (8 participants with CRPS).

### 2.1. Questionnaire measures

All participants completed the Edinburgh Handedness Inventory (EHI; Oldfield, 1971) on which negative scores (< −40) indicate left-handedness and positive scores (> 40), right-handedness. Mood can affect the pain experience (e.g. Tang et al., 2008) and performance on attentional tasks (e.g. Moriya & Nittono, 2011). Therefore, all participants also completed the Profile of Mood States (POMS; McNair, Lorr, & Droppleman, 1971). The Bath CRPS Body Perception Disturbance Scale (BPDS; Lewis & McCabe, 2010) was used to assess subjective cognitive representation of the CRPS-affected / matched limb. This was completed by all participants because it is a non-validated scale with no normative data currently available. Participants with CRPS answered additional questionnaires that were not completed by the control participants. They answered the EHI a second time to rate their *recalled* handedness prior to the onset of CRPS symptoms. An absolute difference between the current and recalled handedness scores (∆EHI) was calculated to approximate the functional impact of the disorder. Pain severity and interference were assessed using a short-form of the Brief Pain Inventory (BPI; Cleeland, 1996) and the neuropathic component of pain was measured using the Pain Detect Questionnaire (Freynhagen et al., 2006). Participants with CRPS also completed the Tampa Scale for Kinesiophobia (Miller et al., 1991), which measures pain-related fear of movement and re-injury. Higher scores on the abovementioned questionnaires indicate greater mood disturbance (POMS), more severe distortion of body representation (BPDS), greater pain severity and interference (BPI), greater neuropathic component of pain (Pain Detect), and more severe kinesiophobia (Tampa Scale).

### 2.2. Sensory, motor, and autonomic function

We used a validated protocol to confirm that participants met the CRPS research diagnostic criteria, to quantify CRPS severity (Harden et al., 2017), and to confirm that the control participants did not present with signs or symptoms of CRPS on their matched limb. We objectively quantified the CRPS signs described below. Temperature asymmetry was quantified as a difference between an average of three hand temperature measurements on the unaffected and affected side (in the centre of the most painful site, and on the dorsal and palmar hand surface over the thenar muscle) using an infrared thermometer (Duratool, thermal resolution 0.1°C). We quantified oedema as a difference between an average of three hands size measurements on the affected and unaffected side using the figure-of-eight procedure (Pellecchia, 2003) with a soft tape measure (cm). Weakness was quantified as a ratio of grip strength in the affected to the unaffected hand, measured as an average of three maximum strength grips of an electronic dynamometer with each hand (kg force; Constant, model 14192-709E). We quantified active range of movement as a ratio of delta finger-to-palm distance (cm; delta refers to the difference between full extension and full flexion of fingers; Torok et al., 2010) in the affected hand to the unaffected hand.

Hypoesthesia, hyperalgesia, and allodynia were additionally quantified using elements of a standardized Quantitative Sensory Testing protocol (Rolke et al., 2006), administered to the centre of most painful region on the affected / matched limb and the corresponding site on the unaffected limb. Mechanical Detection Thresholds were assessed with von Frey filaments (0.008-300g force; Bioseb, model Bio-VF-M). A positive ratio of thresholds for affected vs. unaffected side indicates hypoesthesia (i.e. increased tactile detection threshold) on the affected limb. We used pinprick stimulators (8mN-512mN; MRC Systems Pin Prick Stimulator Set) to quantify Mechanical Pain Thresholds. A positive thresholds ratio for affected vs. unaffected side indicates hyperalgesia (i.e. decreased pain threshold) on the affected limb. Allodynia was assessed by applying with a single sweeping motion a cotton ball, Q-tip, and a brush to the skin, five times each in a random order. Allodynia was quantified as an arithmetic mean of 15 ratings for each sensation from 0 (“no pain, no sharp, pricking, stinging, or burning sensation”) to 100 (“most intense pain sensation imaginable”) on the affected limb. This procedure was adapted from the Dynamical Mechanical Allodynia test (Rolke et al., 2006). We also examined tactile discrimination thresholds on index fingertips of each hand using a Two-Point Discriminator disk (Exacta, North Coast Medical; Pleger et al., 2006). Using a staircase procedure, the participant’s finger was touched with either one tip or two tips of the disk, starting with two points separated by 7mm distance and then increasing or decreasing the distance (down to a single tip) across trials depending on participant’s responses (i.e. whether they reported feeling one or two tips, respectively). The thresholds were calculated as a geometric mean of five subthreshold and five suprathreshold values. A positive thresholds ratio for the affected vs. unaffected side indicates decreased precision of tactile discrimination ability of the affected limb.

### 2.3. Experimental tests of neuropsychological changes

Participants completed three experimental tests of visuospatial attention (the TOJ, Landmark, and Greyscales tasks; see Figure 1a-c), one test of the mental representation of space (the MNLB task; see Figure 1d), and one test of spatially-defined motor function. For convenience, these tasks were completed in the following order: the Landmark task, the Greyscales task, the test of spatially-defined motor function, the TOJ task, and the MNLB. All tasks except the MNLB were administered via PsychoPy software (Peirce, 2007) using a touch-screen laptop computer (Windows 10 operating system, screen dimensions 34.5cm × 19.4cm, resolution 1920 × 1080 pixels). For the tests of visuospatial attention and spatially-defined motor function, the participant’s head was stabilised by a chinrest aligned with a central fixation and positioned at 50cm distance from the screen. Note that the TOJ stimuli were not presented on the computer screen (see section 2.3.2.1.), but participants did use the chinrest. Key-press and key-release responses were recorded using a custom-made button-box. The button-box was aligned with the centre of the screen for all tasks, except for specific blocks of the test of spatially-defined motor function in which it was also placed to the left or right of the screen (see section 2.3.4.). Participants used their unaffected hand to press the buttons in the Landmark and Greyscales tasks, and both hands (one at the time per block) in the test of spatially-defined motor function. When manual responses were not required (i.e. in the TOJ and MNLB tasks), participants rested their uncrossed hands in their lap under the table.

**Figure 1.**
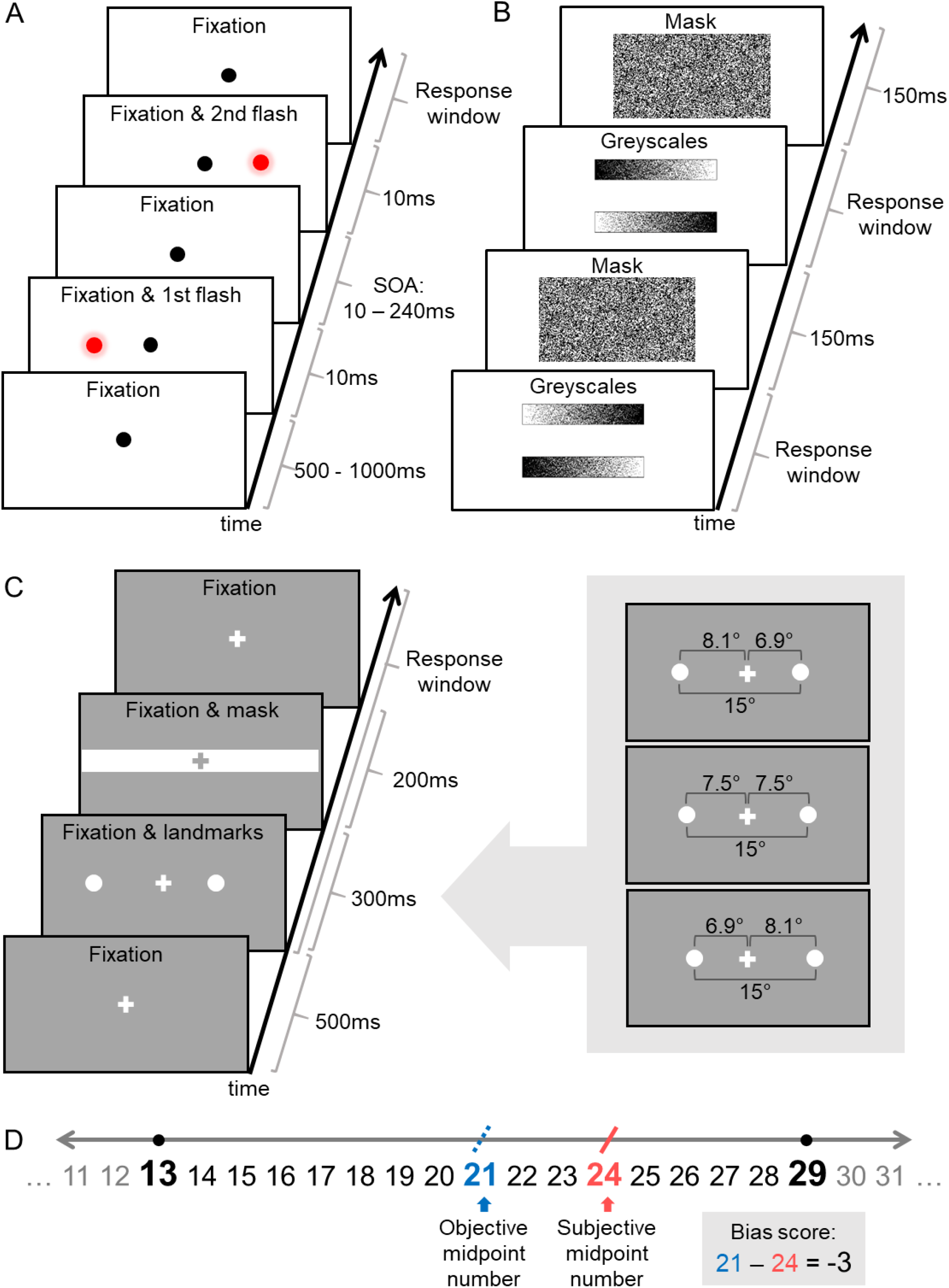
Experimental tests of visuospatial attention (A-C) and the mental representation of space (D). Text within the illustrated screenshots did not appear during the study and is for illustration purposes only. (A) In the Temporal Order Judgement (TOJ) task, the participant maintained their gaze on the central fixation point and verbally reported which light flash (“left” or “right”) appeared first or second, depending on the response block. In each trial, the two lights were presented onto a white table surface with one of ten possible Stimulus Onset Asynchronies (SOAs). (B) In the Greyscales task, the stimuli were on constant display until the participant pressed a button to indicate which of the two greyscale bars (upper or lower) appeared overall darker. Then the stimuli were replaced by a mask, and the next trial began. (C) In the Landmark task, the participant maintained their gaze on the fixation cross and pressed a button to indicate which landmark (left or right) appeared further from or closer to the fixation cross, depending on the response block. The right inset panel illustrates three out of 13 possible arrangements of the landmark stimuli (lines and numbers are for illustration purposes only). The top rectangle corresponds to the left landmark being furthest from, and the right landmark being closest to fixation; the middle rectangle corresponds to both landmarks being equidistant from fixation; and the bottom rectangle corresponds to the left landmark being closest to, and the right landmark begin furthest from fixation. The distance between the two landmarks was constant (15°), while their relative horizontal distance from fixation varied by 0.1° across trials. In each trial, the landmarks were replaced by a mask after 300ms. (D) In the Mental Number Line Bisection task, the experimenter presented each trial verbally, e.g. “What is the midpoint number between 13 and 29?”. The participant verbally reported their subjective midpoint number (e.g. “24”) without making any calculations. Deviation from the objective midpoint (e.g. “21”) on each trial was calculated by subtracting the subjective midpoint from the objective midpoint. In all four tasks, participant’s non-speeded response initiated the next trial.

The data from the computer tasks were transformed to reflect the participants’ performance relative to their affected / unaffected side of the body or visual field. For example, participants with CRPS whose right limb was affected would have their responses to left-sided stimuli coded as “unaffected side”, and responses to right-sided stimuli as “affected side” (and vice versa for participants whose left limb was affected). To enable comparison of both groups relative to affected and unaffected side, control participants’ limbs were coded as “affected” and “unaffected” with respect to their matched participant with CRPS (regardless of the participants’ handedness).

#### 2.3.2. Visuospatial attention

##### 2.3.2.1. The Temporal Order Judgement (TOJ) task

For the TOJ task, participants verbally reported the order of two brief (10ms) identical lights that were projected onto a white table surface using laser pointers controlled via an Arduino platform. The lights appeared one 9cm to the left and one 9cm to the right of a central fixation point located approximately 28cm away from participant’s torso. Pairs of lights were presented 15 times for each of the ten temporal offsets (±10, ±30, ±60, ±120 and ±240ms; negative values represent the trials in which the left light appeared first) in pseudorandom order, resulting in 150 trials (Figure 1a). To account for potential response biases (Filbrich et al., 2016), participants completed the TOJ task twice: once indicating which light appeared first, and once indicating which light appeared second (block order was counterbalanced). For each block, the relative number of participant’s “right light appeared first” or “left light appeared second” responses to the range of temporal offsets was fitted with a cumulative Gaussian using a criterion of maximum likelihood to obtain the Point of Subjective Simultaneity (PSS). Following the transformation from left / right to affected / unaffected, the PSS values were averaged between the two response blocks to give a single value for each participant. The PSS is an index of spatial attention bias and indicates the amount of time (ms) by which the light on the affected side must precede (negative PSS) or follow (positive PSS) the light on the unaffected side for the two stimuli to be perceived as simultaneous. Therefore, negative PSS values indicate reduced attention to the affected relative to the unaffected side and positive numbers indicate greater attention to the affected relative to the unaffected side.

##### 2.3.2.2. The Landmark task

We designed a version of the Landmark task based on one previously used to demonstrate underrepresentation of the side of near space corresponding to the missing limb in amputees (Makin et al., 2010). In each trial, two identical landmarks (white circles) were simultaneously presented on a computer screen with a fixed distance between them (15°), but in different positions relative to central fixation. The stimulus locations varied from ±8.1° to ±6.9° away from fixation in the horizontal plane by 0.1° increments (e.g. −8.1° and +6.9°, −8.0° and +7.0°, −7.9° and +7.1°, etc., up to −6.9° and +8.1°; see Figure 1c for example stimulus pairs). Negative values represent the location of the left landmark, and positive values the location of the right landmark, with reference to central fixation at 0°. Each pair of landmarks was presented 15 times in each of 13 possible arrangements (including equidistant), resulting in 195 trials. Similar to the TOJ task, participants completed the Landmark task twice to account for any response biases: once indicating which landmark appeared further from fixation, and once indicating which landmark appeared closer to fixation (block order was counterbalanced). For each block, the relative number of participant’s keypress responses to the range of spatial offsets indicating “right landmark appeared further” or “left landmark appeared closer” was fitted with a cumulative Gaussian to derive the Point of Subjective Equality (PSE). Following the transformation from left / right to affected / unaffected, the PSE values were averaged between the two response blocks to give a single value for each participant. The PSE is an index of spatial bias that represents the relative distance (°) at which the landmark on the affected side should be further from (negative PSE) or closer to (positive PSE) central fixation for the two landmarks to be perceived as equidistant. Therefore, negative PSE values indicate under-representation of the affected side of space relative to the unaffected side and positive values indicate over-representation of the affected side of space relative to the unaffected side.

##### 2.3.2.3. The Greyscales task

In each trial of the Greyscales task (Nicholls et al., 1999), participants were presented with two vertically aligned greyscale bars that were positioned one on top of the other. Each bar was darker at one end than the other, and the two bars were mirror images of each other such that one was darker on the left and the other was darker on the right even though both bars had the same average luminance (Figure 1b). Participants indicated with a button press which bar (top or bottom) was darker overall (in free-viewing conditions). The number of times the participant chose a bar that was darker on its right side, regardless of its vertical position, was subtracted from the number of times the participant chose a bar that was darker on its left side. We then divided this value by total number of trials (i.e. 40) to calculate an index of spatial attention bias. Transformed negative scores indicate reduced attention to the affected side of space, consistent with making higher proportion of relative darkness judgements based on the side of the stimuli corresponding to the unaffected limb.

#### 2.3.3. Mental representation of space

We used a Mental Number Line Bisection (MNLB) task based on that of Sumitani et al. (2014). In each trial, participants were instructed to verbally estimate, without calculating, the midpoint number between a given pair of numbers (Figure 1d). There were 84 trials with pairs of numbers separated by intervals of 9, 16, 25, 39, 49, and 64 digits, with the individual numbers ranging from 2 to 98. Number pairs were read aloud by the researcher in pseudorandom order. To account for potential response bias, each numbers pair was presented once in ascending (e.g., 54 and 70) and once in descending (e.g., 70 and 54) order. Individual spatial bias scores were computed by subtracting participant’s subjective midpoint number from the objective midpoint number in each trial and averaging the results across trials. A negative index indicates a relative bias towards guessing larger numbers as the midpoint number. That is, following the transformation from left (smaller numbers) / right (larger numbers) to affected / unaffected, a negative index indicates a bias away from the affected side of the mental representation of space.

#### 2.3.4. Spatially-defined motor function

We adapted a test for directional hypokinesia previously used in research on hemispatial neglect (Sapir et al., 2007) to test for spatially-defined (directional) motor deficits in CRPS patients. Each trial was initiated by the participant holding down a button with an index finger, while maintaining their gaze on a central fixation cross flanked by two squares located 12° to the left (left Visual Field, VF) and 12° to the right (right VF) from fixation. After a time interval that randomly varied between 1500ms and 3000ms, a target (“X”) appeared in one of the squares for 2000ms. The target location was pseudorandomized across 30 trials within each block and expressed relative to the CRPS-affected / matched side (i.e. in terms of the affected and unaffected VFs rather than left and right VFs). Participants were instructed to make speeded movements to release the button and touch the target location on the touch-screen using the same finger, and then return their hand to hold down the button, which initiated the next trial. We recorded the reaction time to release the button after target onset (movement initiation time) and the time between releasing the button and touching the screen (movement execution time). There were three hand Starting Positions in which the button box was either aligned with the body midline, or located 25cm to the left or to the right from the body midline. These locations were expressed relative to the CRPS-affected (or matched “affected”) limb, that is, as the central, affected, and unaffected Starting Positions. Participants completed six blocks of the task in total: two blocks from each Starting Position (order counterbalanced), one with each Hand (alternating between the affected and unaffected hand between consecutive blocks). Slower initiation and execution of movements directed towards the affected side of space, independent of the hand used, would be taken to indicate directional hypokinesia and bradykinesia, respectively.

### 2.4. Data handling and statistical analyses

The data was processed and analysed using MATLAB 2018b, IBM SPSS Statistics 25, R 3.5.3, and JASP 0.9.2.0 software. The significance level for frequentist hypotheses testing was *α* = .05. For Bayesian analyses we used the suggested cut-offs for Bayes factor (*BF*_*10*_; Lee & Wagenmakers, 2014). We used Holm-Bonferroni correction for multiple comparisons to control for family-wise type I error in the primary analyses. Corrections were not implemented in exploratory analyses.

Pre-processing of the data from the spatially-defined motor function task involved removing invalid trials from individual data sets, i.e. trials in which the screen touch did not match the target location or the button was released before target onset (for movement initiation time analysis), and additionally the trials in which screen touch time was not recorded (for movement execution time analysis). In total, 7.25% of all completed trials were removed across all participants. Outliers in participant-level data in this task were identified as scores outside ±3 *SDs* from the participant’s score for a task condition, and replaced with the nearest non-outlier values (0.84% of all valid data replaced). Missing questionnaire items in participant-level data were replaced with participant’s mean rating for the specific subscale calculated without the missing items (person mean replacement; 0.08% of all questionnaire items replaced across all participants).

For group-level data, scores outside ±3 *SDs* from the group mean for each test or task condition were identified as outliers and replaced with the nearest non-outlier values. Missing data points on clinical measures and computer-based tasks were replaced with a group mean for particular test or task condition, with the exception of the test of spatially-defined motor function (six participants with CRPS could only complete the task using the unaffected limb, thus they were excluded from the affected limb analysis). In group-level data, 1.22% of all data points were replaced as missing or outlying values across all measures and all participants.

Bootstrapping was implemented for descriptive and inferential statistics using 1000 bootstrap samples and calculating bias corrected and accelerated 95% confidence intervals (BCa 95% CIs). For non-parametric tests, we used Monte Carlo estimation of 95% CIs based on 1000 samples. Between-group differences on categorical variables were estimated through chi-square statistics. To compare mean scores on the continuous variables between participants with CRPS and control participants, we conducted *t* tests and ANOVAs, and interrogated significant interactions through contrasts. Where assumptions of *t-*tests were violated, we carried out Wilcoxon signed-rank tests and Mann-Whitney *U* tests and reported median scores. Due to missing data and violations of normality, homogeneity of variance, and sphericity assumptions for ANOVAs in the data from the test of spatially-defined motor function, bootstrapped linear mixed models analyses were conducted instead to investigate the interactions of interest. To investigate any potential relationships between neuropsychological changes and clinical signs of CRPS in the data from participants with CRPS, we conducted exploratory best subsets regression analyses.

## 3. Results

### 3.1. Participant characteristics; questionnaire measures; and sensory, motor, and autonomic function

Group-level participant characteristics are reported in Table 1, including average scores on the self-reported measures of pain, kinesiophobia, body perception disturbance, mood, and hand preference; and tests of sensory, motor, and autonomic function. Participants with CRPS and controls were equally matched on mean age, proportion of males and females, and proportion of left- and right-handed participants in each group (*p*s > .05). Comorbidities and current treatments of participants with CRPS are summarised in the Supplementary Material.

**Table 1.**
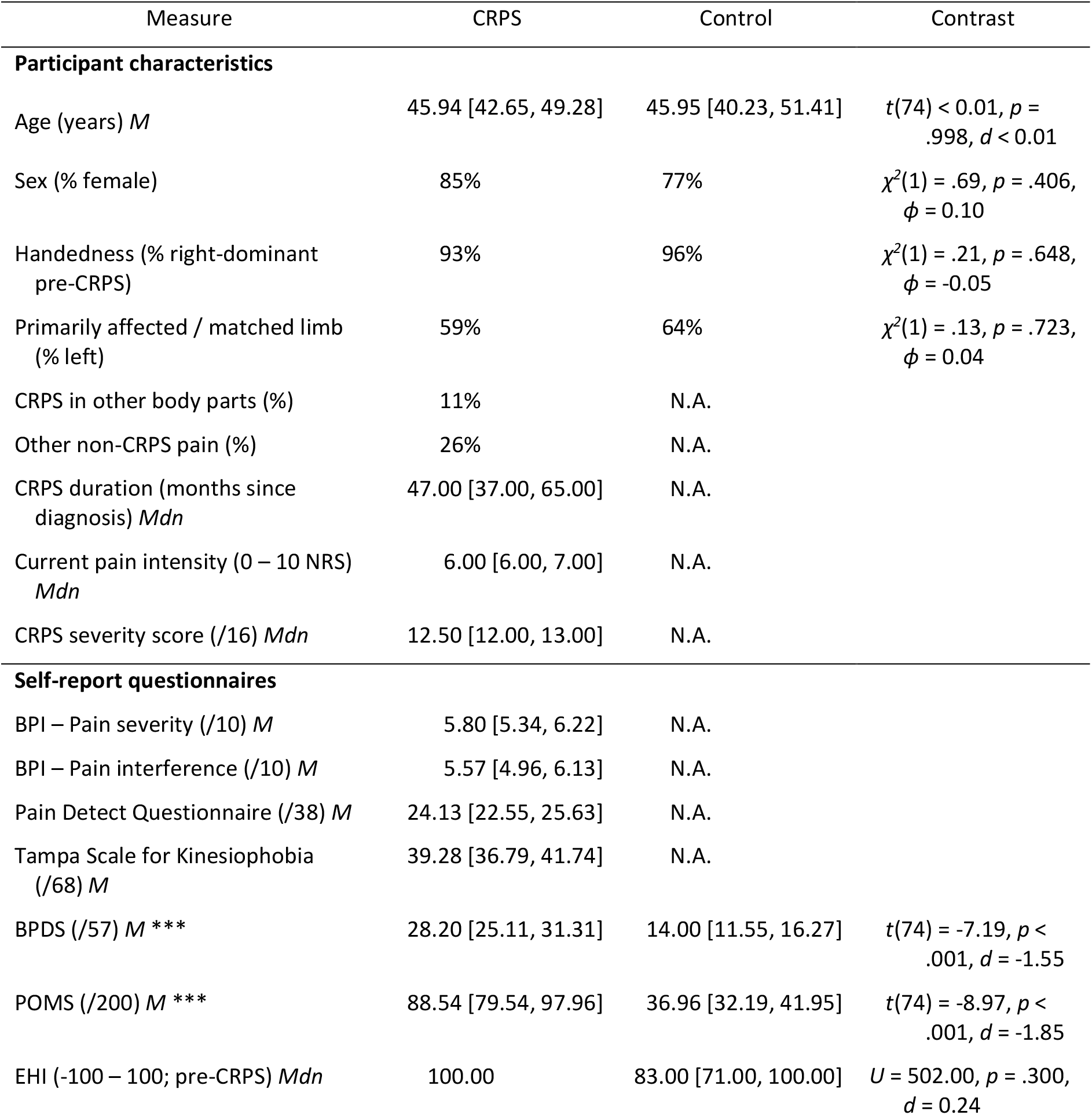
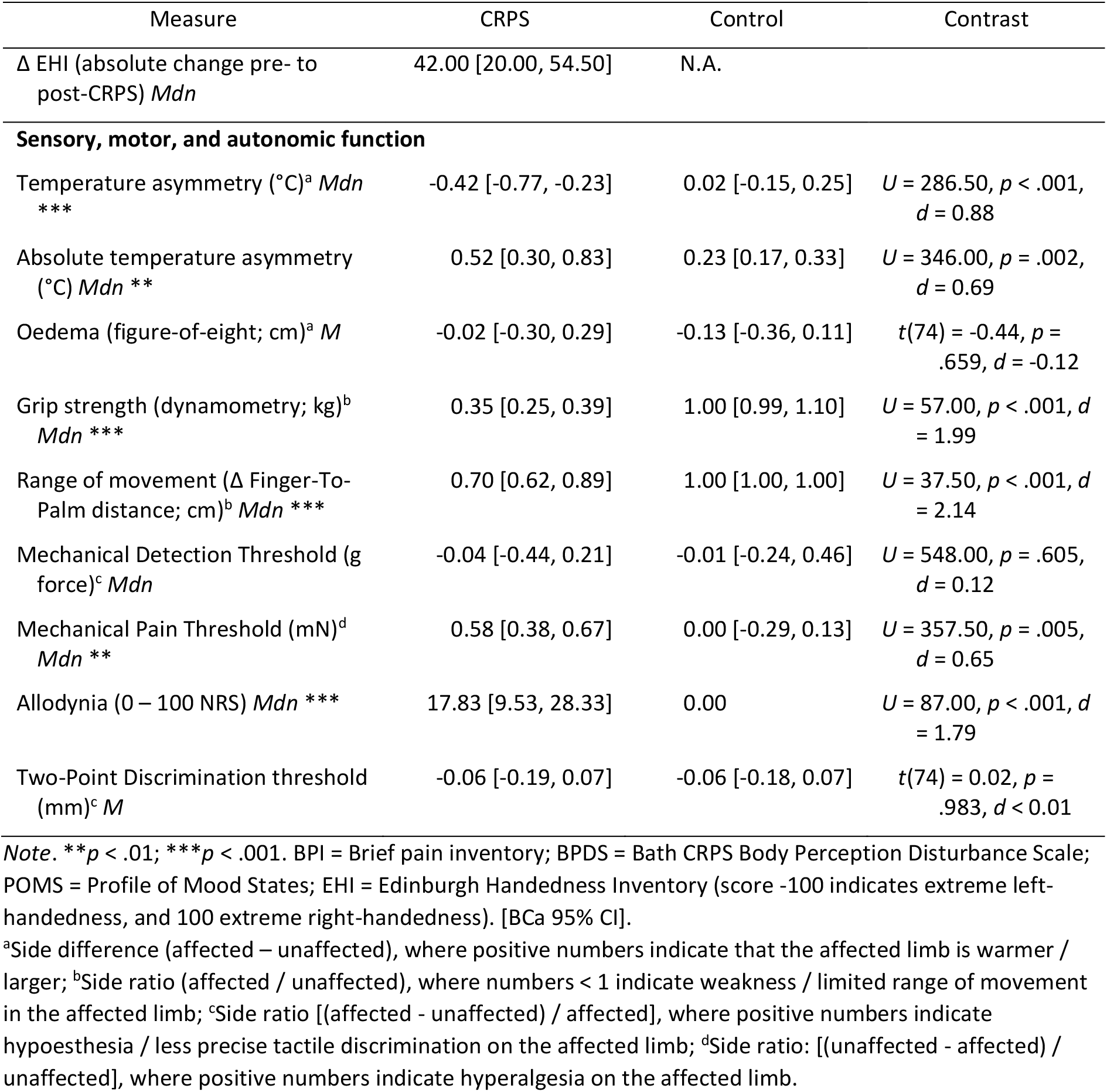
Group-level participant characteristics; scores on questionnaire measures; and quantification of sensory, motor, and autonomic function of the affected limb relative to the unaffected limb in participants with CRPS compared to healthy controls

Participants with CRPS reported moderate pain severity and interference on the BPI (Li et al., 2007) and their mean score on the Pain Detect Questionnaire (≥ 19 cut-off) suggested a likely neuropathic pain component (Freynhagen et al., 2006). Despite comparable pain intensity, median CRPS severity score in our sample was higher than in a group of people with chronic (on average 35 months) CRPS tested in the severity score validation study (Harden et al., 2017). This could be because we only included people who met more stringent Budapest research (compared to clinical) diagnostic criteria. The mean score on the Tampa Scale for Kinesiophobia indicated high pain-related fear of movement, comparable with previous CRPS research (Velzen et al., 2019). BPDS scores of participants with CRPS were significantly higher compared to controls, indicating significantly distorted perception of the CRPS-affected limb. POMS scores were also significantly higher among participants with CRPS than controls, indicating greater mood disturbance. There was no difference between the median handedness index of the control participants and recalled (pre-CRPS) handedness of participants with CRPS. Sixty-nine percent of those whose dominant arm was affected by CRPS, or who were ambidextrous before the symptoms onset, showed a change in hand preference towards the unaffected arm according to the absolute difference in the scores on the EHI answered with regard to current and pre-CRPS handedness.

Participants with CRPS presented with significantly larger asymmetries between the affected and unaffected limbs compared to controls in limb temperature (both signed and absolute difference), grip strength, finger-to-palm distance, and mechanical pain threshold (see Table 1). Specifically, the affected limb was on average characterised by lower temperature, weaker grip strength, more limited range of movement, and greater hyperalgesia. Participants with CRPS also had significantly more severe allodynia on the affected limb than the control participants. There were no significant between-group differences in oedema, mechanical detection thresholds, and two-point discrimination thresholds.

### 3.4. Experimental tests of neuropsychological changes

#### 3.4.2. Visuospatial attention

Two-tailed contrasts showed that the performance of participants with CRPS on the three tasks measuring visuospatial attention did not significantly differ from the performance of healthy controls. Specifically, the PSS values in the visual TOJ task were not significantly different between the participants with CRPS (*Mdn* = −0.27; BCa 95% CI [−7.61, 3.97]) and controls (*Mdn* = −5.17, BCa 95% CI [−10.97, 0.27]), *U* = 472.00, *p* = .152, *d* = 0.33 (Figure 2a). Similarly, the PSEs of participants with CRPS in the Landmark task (*Mdn* = 0.05; BCa 95% CI [−0.04, 0.10]) did not significantly differ from the PSEs of the control participants (*Mdn* = 0.06, BCa 95% CI [−0.15, 0.26]), *U* = 551.00, *p* = .624, *d* = 0.11 (Figure 2b). Finally, there were no between-group differences in the bias scores on the Greyscales task (CRPS: *M* = 0.11, BCa 95% CI [−0.02, 0.23]; controls: *M* = 0.01, BCa 95% CI [−0.17, 0.19]), *t*(74) = −0.81, *p* = .422, *d* = −0.20; Figure 2c). Follow up Bayesian analyses using a Cauchy prior width of 0.707 indicated anecdotal evidence of no difference between groups for PSSs (*BF*_*10*_ = 0.44) and Greyscales bias scores (*BF*_*10*_ = 0.34), and moderate evidence of no difference between groups for PSEs (*BF*_*10*_ = 0.27) (Lee & Wagenmakers, 2014). CRPS participants’ performance on the TOJ task did not correlate with the other visuospatial tasks, but there was a moderate positive relationship between their scores on the Greyscale and Landmark tasks (*r* = 0.40; see Supplementary Figure 2).

**Figure 2.**
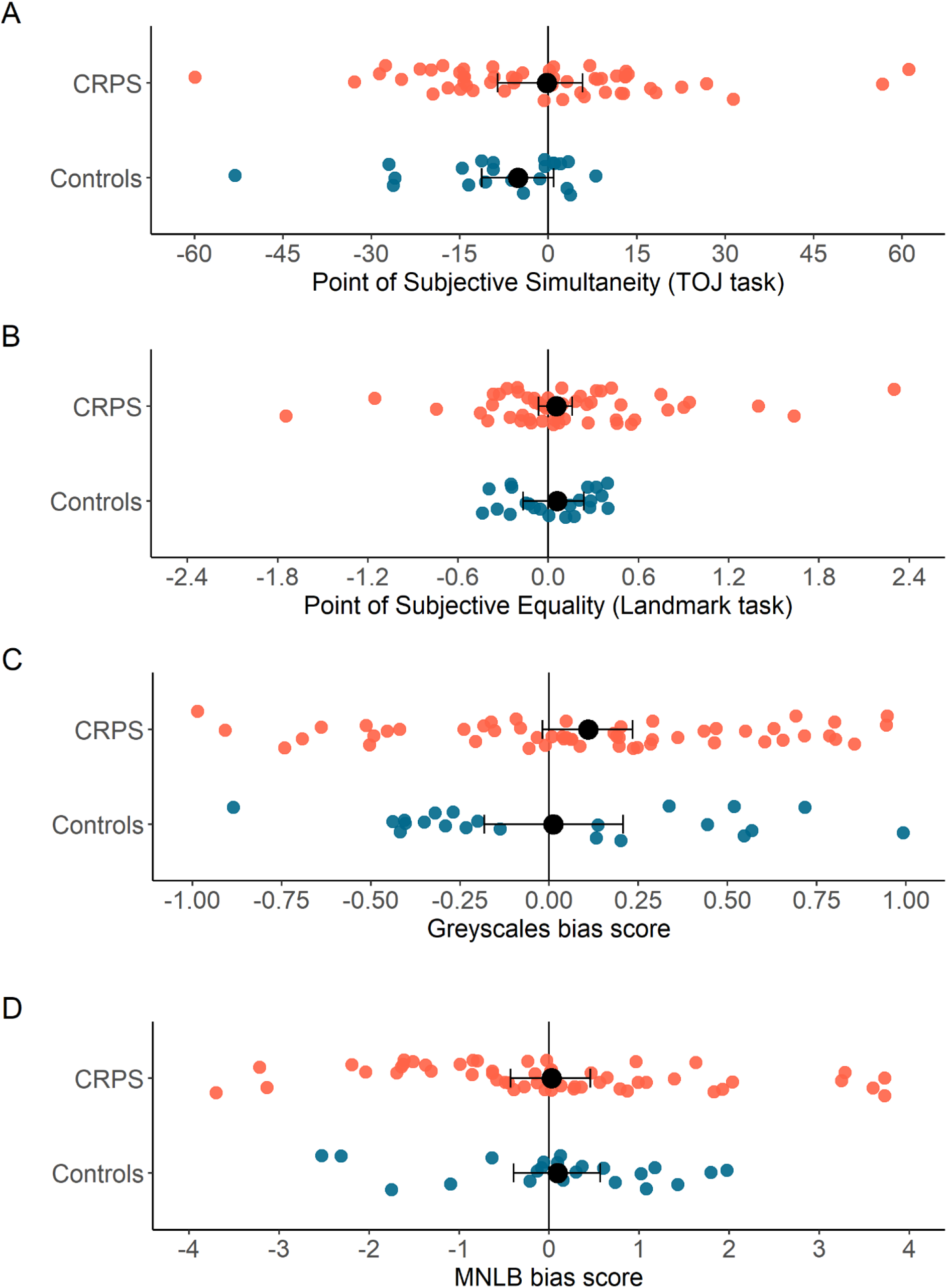
Results of the experimental tests of visuospatial attention (A-C) and the mental representation of space (D). Smaller circles represent individual data from participants with CRPS (orange) and pain-free control participants (blue). Larger black circles represent the group median (A, B) and mean (C, D) scores with bootstrapped 95% confidence intervals (error bars). (A) The Point of Subjective Simultaneity on the Temporal Order Judgement task indicates by how many milliseconds the light on the affected side should precede (negative values) or follow (positive values) the light on the unaffected side for the two lights to be perceived as simultaneous. (B) The Point of Subjective Equality on the Landmark task indicates by how many degrees of visual angle the pair of landmarks should be offset from being truly equidistant to central fixation towards the affected side (negative values) or towards the unaffected side (positive values) for the two landmarks to be perceived as equidistant. (C) The bias score on the Greyscales task indicates to what extent the participants were basing their darkness judgements on the side of the stimuli corresponding to their unaffected side (negative values) or to their affected side (positive values). (D) The bias score on the Mental Number Line Bisection task indicates to what extent participants’ subjective midpoint of the mental number line was shifted towards the numbers corresponding to their unaffected side (e.g. higher numbers for participants with left-CRPS; negative values) or to their affected side (e.g. smaller numbers for participants with left-CRPS; positive values). Negative scores for each of the measures depicted in this figure would indicate reduced attention to or (mental) representation of the affected side of space.

#### 3.4.3. Mental representation of space

MNLB bias scores of the participants with CRPS (*M* = 0.02, BCa 95% CI [−0.39, 0.43]) were not significantly different to those of the healthy controls (*M* = 0.09, BCa 95% CI [−0.52, 0.65]; *t*(74) = 0.18, *p* = .860, *d* = 0.05; Figure 2d), suggesting unbiased mental representation of space. A Bayesian independent-samples *t*-test indicated moderate evidence of no difference (*BF*_*10*_ = 0.26). CRPS participants’ performance on the MNLB task was moderately positively correlated with their performance on the Greyscales and Landmark tasks (*r* = 0.37 and 0.38; see Supplementary Figure 2).

#### 3.4.4. Spatially-defined motor function

##### 3.4.4.1. Linear mixed models regressions on movement initiation and execution times

After excluding incorrect and missed trials, we computed mean movement initiation time and movement execution time for each combination of VF and hand Starting Position. The tasks performed with the affected (CRPS *n* = 43 [initiation], 45 [execution]; control *n* = 21) and unaffected limb (CRPS *n* = 50; control *n* = 18) were analysed separately.

The data for this task were analysed using four bootstrapped linear mixed models regression analyses. The four outcome measures were initiation times and execution times for both the affected and unaffected limb. The fixed effects for each analysis were Group (participants with CRPS, healthy controls), Starting Position (affected, central, unaffected), VF (affected, unaffected) and their interactions. Participant ID was entered as a random effect in each analysis. As this method is robust to the presence of outliers and missing values (Wu, 2009), we used unprocessed data (i.e. data prior to replacement of group-level outliers and missing values). A variable was considered to be making a significant contribution to predicting the outcome variable when the 95% CI around the regression coefficient (B) did not include zero. As our main objective was to assess the differences in motor function between participants with CRPS and healthy controls, here we only summarise those significant main effects and interactions that involved Group. The full results for all four regression analyses are reported in Supplementary Material.

The terms of the regression analyses that were of most interest in the present study were the interactions between Group and VF; and between Group, Starting Position, and VF. The results showed that these terms did not significantly contribute to the prediction of movement initiation and execution times for either the affected or unaffected hand (i.e. all confidence intervals around the relevant regression coefficients included 0; see Supplementary Table 1).

We found significant main effects of Group on initiation times for both limbs (affected *B* = 0.19, BCa 95% CI [0.16, 0.22]; unaffected *B* = 0.07, BCa 95% CI [0.05, 0.10]), indicating that, regardless of the hand Starting Position or the VF in which the target appeared, participants with CRPS were slower to initiate movements with their affected (*Mdn* = 553.28, BCa 95% CI [488.40, 582.85]) and unaffected (*Mdn* = 459.50, BCa 95% CI [438.19, 485.04]) limbs compared to the initiation times of the control participants with their matched “affected” (*Mdn* = 416.09, BCa 95% CI [403.52, 435.54]) and “unaffected” (*Mdn* = 412.74, BCa 95% CI [394.47, 438.38]) limbs. The analyses of movement execution times also showed significant main effects of Group for both limbs (affected *B* = 0.44, BCa 95% CI [0.36, 0.52]; unaffected *B* = 0.11, BCa 95% CI [0.08, 0.15]). Specifically, execution of movement with the affected (*Mdn* = 970.45, BCa 95% CI [907.66, 1012.56]) and unaffected (*Mdn* = 820.14, BCa 95% CI [733.29, 858.03]) limbs among participants with CRPS was slower compared to execution times with the matched “affected” (*Mdn* = 677.37, BCa 95% CI [620.88, 746.19]) and “unaffected” (*Mdn* = 678.87, BCa 95% CI [586.43, 756.35]) limbs in the control group. These effects are consistent with overall slowing of initiation and execution of movements with both affected and unaffected limbs in participants with CRPS relative to healthy controls.

The regression model for movement execution times with the unaffected limb showed that the term for the interaction between Group and affected versus unaffected Starting Position was significant (*B* = 0.06, BCa 95% CI [0.002, 0.12]). For participants with CRPS, the difference in execution times for movements originating from the unaffected (*Mdn* = 858.14, BCa 95% CI [795.38, 887.98]) compared to affected (*Mdn* = 801.83, BCa 95% CI [722.31, 811.46]) Starting Positions was larger relative to the same difference for controls (unaffected *Mdn* = 692.69, BCa 95% CI [643.87, 769.59]; affected *Mdn* = 658.19, BCa 95% CI [590.84, 771.66]), regardless of the VF in which the targets appeared (Figure 3). This pattern is consistent with directional bradykinesia for the affected space: slowing of movements directed toward the affected side of space relative to movements directed toward the unaffected side of space. In the same regression model, the term for the Group by affected versus central Starting Position interaction was not significant (*B* = 0.01, BCa 95% CI [−0.04, 0.06]).

**Figure 3.**
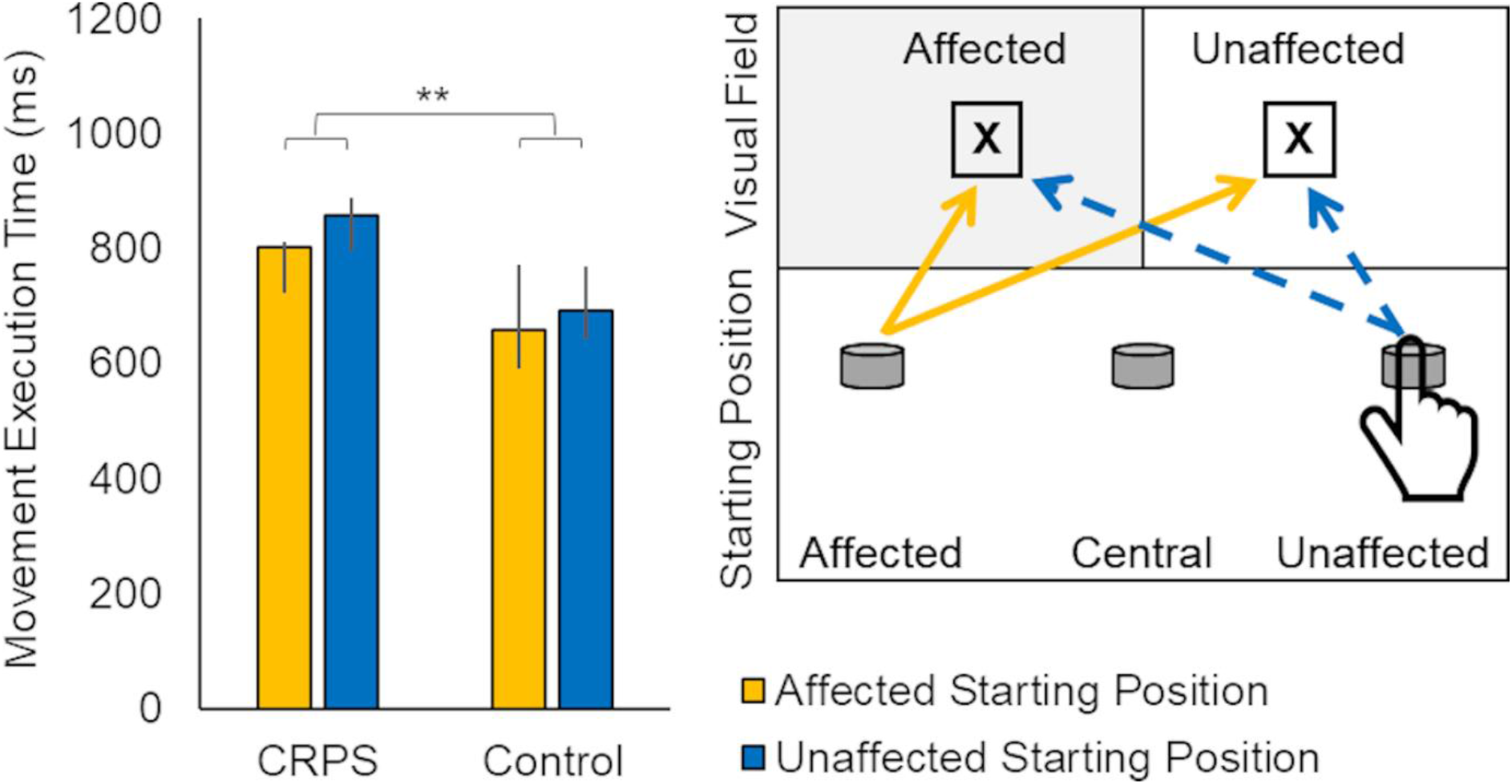
Interaction between Group and Starting Position on execution times of movements performed with the unaffected limb starting from the affected compared to unaffected positions. Bars represent CRPS and control participants’ median execution times (error bars: BCa 95% CIs) with the unaffected hand from affected (yellow) and unaffected (blue) Starting Positions, averaged across two Visual Fields. **The interaction is significant at the level of *p*_adjusted_ < .01. The inset panel (right) illustrates slower execution of movements to the targets (X) in either Visual Field from the unaffected Starting Position (blue dashed arrows), relative to the affected Starting Position (yellow solid arrows).

##### 3.4.4.2. Analyses of directional hypokinesia and bradykinesia indices

To dissociate any signs of directional hypokinesia and bradykinesia from potential visual “neglect-like” deficits, biomechanical constraints, and lengths of movement pathways from different starting positions, we additionally analysed specific indices for each limb separately. We calculated two indices of directional hypokinesia towards the affected side based on those used in previous research on spatial motor biases in stroke patients (Sapir et al., 2007). The relevant movement pathways and formulae are represented in Figure 4. The first index (A; Figure 4a) describes the difference in initiation times towards the affected VF with respect to the unaffected VF, depending on the direction of movements (that is, as a function of starting position). We derived a second index (B; Figure 4b), which in contrast to Index A, does not involve comparing a movement within one side of space to one across the body midline (and therefore over a longer pathway). Index B directly describes the relative slowing (if any) of initiations of movements to the affected VF when making movements of the same physical length directed toward the affected side compared to movements directed toward the unaffected side (Figure 4d). Larger (more positive) values of Indices A and B indicate greater directional hypokinesia towards the affected side. To account for the possibility of directional hypokinesia towards the *unaffected* side (i.e. in the direction opposite to hypothesized “neglect-like” motor deficits), we computed two additional indices (C and D; Figure 4c,d), which were not considered in Sapir et al.’s (2007) study. Indices C and D are analogous to Indices A and B, respectively, and describe relative slowing of movement initiation toward the unaffected side with respect to the affected side. Larger values of Indices C and D indicate greater directional hypokinesia towards the unaffected side. We calculated the same four indices for movement execution times to examine any signs of directional bradykinesia. We examined differences between participants with CRPS and healthy controls on each index and for each hand through separate between-group contrasts.

**Figure 4.**
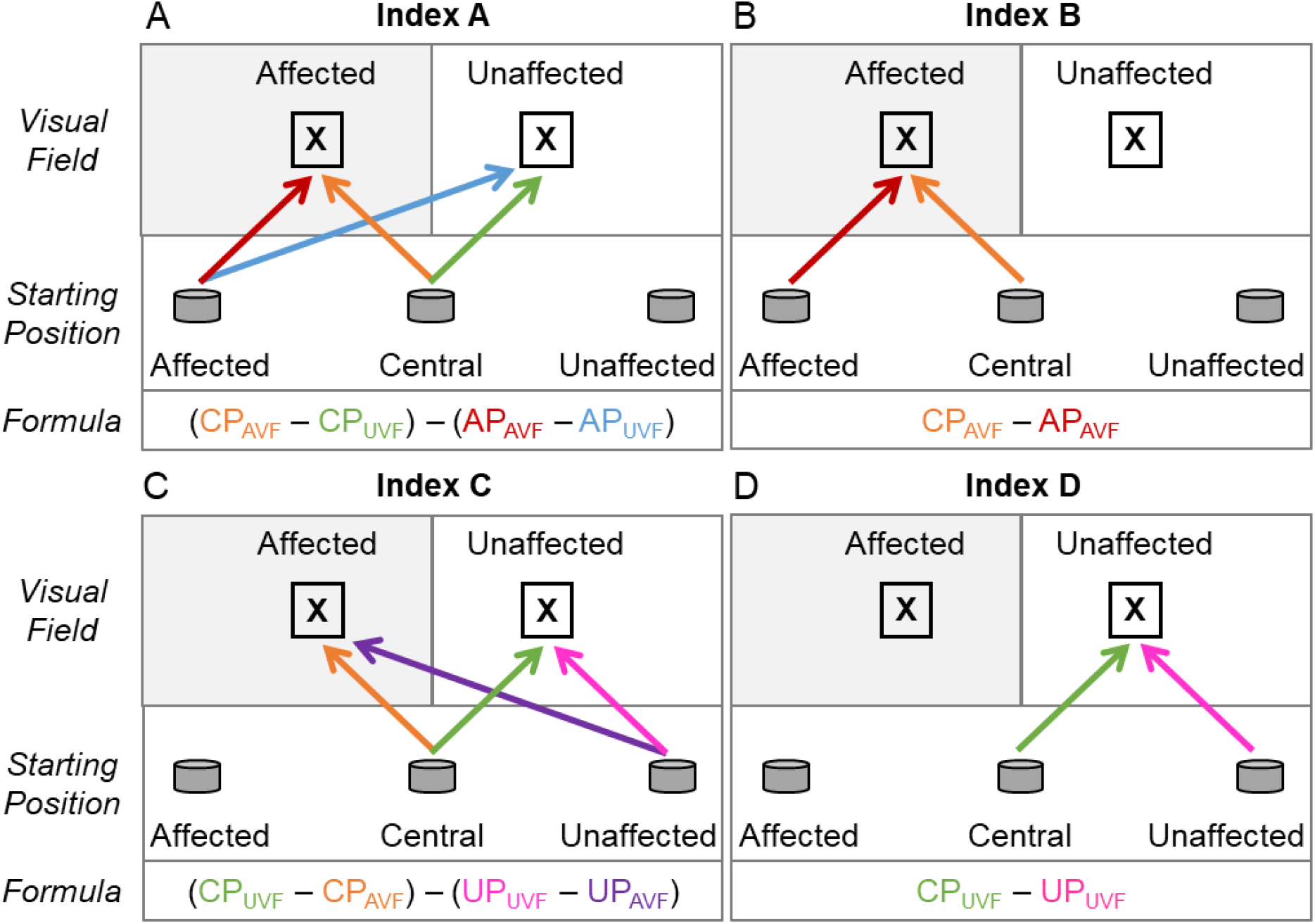
Movement pathways and formulae used to calculate indices of directional hypokinesia and bradykinesia towards the affected (A, B) and unaffected (C, D) side of space. The arrows indicate the direction of movement from hand Starting Position (affected, AP; central, CP; and unaffected, UP) to the targets (X) appearing in the affected (AVF) or unaffected (UVF) Visual Field. Indices for each hand were computed using movement initiation (hypokinesia) or execution (bradykinesia) times according to the formulae represented in the bottom segments of the each panel. More positive values for Indices A (A) and B (B) would indicate greater directional hypokinesia/bradykinesia towards the affected side; more positive values for Indices C (C) and D (D) would indicate greater directional hypokinesia/bradykinesia towards the unaffected side.

After Holm-Bonferroni correction, Mann-Whitney *U* tests did not show significant differences between the CRPS participants’ and controls’ indices of directional hypokinesia and bradykinesia towards the affected side (*Us* ≥ 323.00, *ps*_*adjusted*_ ≥ .062, *ds* ≤ 0.47), with one exception. Index A (Figure 4a) for movement execution with the unaffected limb was significantly more positive among the CRPS participants (*Mdn* = 100.16, BCa 95% CI [84.22, 125.09]) compared to control participants (*Mdn* = 59.42, BCa 95% CI [39.37, 73.29]), *U* = 294.00, *p*_*adjusted*_ = .032, *d* = 0.55. Analysis of the indices of directional hypokinesia and bradykinesia towards the unaffected side showed that the CRPS participants’ Index C (Figure 4c) for movement initiation with the affected limb (*Mdn* = 18.05, BCa 95% CI [3.02, 30.89]) was significantly more positive than the controls’ Index C (*Mdn* = 0.19, BCa 95% CI [−13.01, 8.25]), *U* = 272.00, *p*_*adjusted*_ = .016, *d* = 0.63. There were no other significant between-group differences (*Us* ≥ 318.00, *ps*_*adjusted*_ ≥ .072, *ds* ≤ 0.46). Overall, these results indicate that there was some evidence for participants with CRPS showing significant directional bradykinesia towards the affected side (Index A) when using the unaffected limb, but also for significant directional hypokinesia towards the unaffected side (Index C) when using the affected hand, compared to controls.

Considering that only a subset of stroke patients in Sapir et al.’s (2007) study presented with significant directional hypokinesia (9 out of 52 patients, i.e. 17%, in a task performed only with the unaffected hand; identified based on z-scores compared to controls’ distribution), we explored whether there was a subgroup of CRPS patients showing this deficit. For this purpose, we compared each individual patient’s Indices A and B for movement initiation and execution with the affected and unaffected hand to the controls’ mean indices using Crawford *t*-tests (Crawford & Howell, 1998). A patient was classified as showing signs of directional hypokinesia or bradykinesia towards the affected side if both their Indices (A and B) were significantly more positive than controls’ mean indices (*p*s < .05). For balance, we used the same method to explore what proportion of patients presented with significant directional hypokinesia or bradykinesia towards the *unaffected* side, that is, had more positive Indices C and D. Table 2 summarises the results. Overall, when the affected limb was used, directional hypokinesia and bradykinesia towards the unaffected side was more prevalent than towards the affected side, and the opposite tendency was seen when the unaffected limb was used. However, the absolute number of patients with signs of directional hypokinesia and bradykinesia when the unaffected limb was used was low.

**Table 2.**
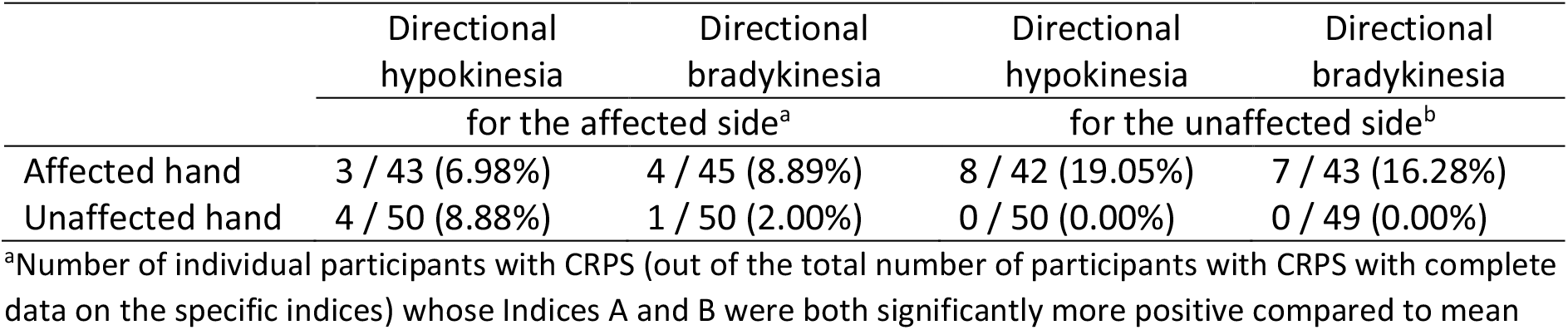
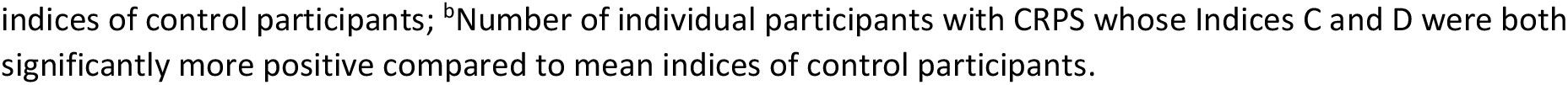
Proportion of participants with CRPS who showed signs of directional hypokinesia or bradykinesia towards the affected side (Indices A and B) or towards the unaffected side (Indices C and D)

|  | Directional<br>hypokinesia<br>for the affected side<sup>a</sup> | Directional<br>bradykinesia<br>for the affected side<sup>a</sup> | Directional<br>hypokinesia<br>for the unaffected side<sup>b</sup> | Directional<br>bradykinesia<br>for the unaffected side<sup>b</sup> |
| --- | --- | --- | --- | --- |
| Affected hand | 3 / 43 (6.98%) | 4 / 45 (8.89%) | 8 / 42 (19.05%) | 7 / 43 (16.28%) |
| Unaffected hand | 4 / 50 (8.88%) | 1 / 50 (2.00%) | 0 / 50 (0.00%) | 0 / 49 (0.00%) |

<sup>a</sup>Number of individual participants with CRPS (out of the total number of participants with CRPS with complete data on the specific indices) whose Indices A and B were both significantly more positive compared to mean
indices of control participants; <sup>b</sup>Number of individual participants with CRPS whose Indices C and D were both significantly more positive compared to mean indices of control participants.

#### 3.4.5. Relationships between neuropsychological changes and clinical symptoms of CRPS

To investigate the relationships between neuropsychological changes and clinical symptoms of CRPS, we conducted best subsets regression analyses on the data from participants with CRPS. Due to the exploratory nature of this analysis, we took this automated approach to avoid any biased selection of the predictors. The pool of potential predictors of each outcome included all measures of neuropsychological, sensory, motor, autonomic, and psychological changes, as measured by computer-based tasks, clinical and sensory assessments, and self-reported questionnaires. Best subsets regressions were determined for the outcome variables BPI pain severity and CRPS severity score, as key measures of clinical severity of this condition. We also performed regressions on those neuropsychological outcomes on which participants with CRPS differed from controls: BPDS, movement initiation time with the affected hand, movement initiation time with the unaffected hand, movement execution time with the affected hand, and movement execution time with the unaffected hand. The only pre-selection involved removing the variables that were not linearly related to the outcome of interest, to satisfy the assumption of linearity. To address co-linearity, when two variables were highly correlated with each other (Pearson’s *r* > .70; see Supplementary Figure 2), only the one with higher correlation with the outcome was entered into regression analysis. This was the case for the following pairs of variables: current pain intensity and BPI pain severity; BPI pain severity and BPI pain interference; movement initiation time of the affected and unaffected hand; and signed and absolute temperature difference. Considering our sample size (N = 54), we compared best subsets regression models that included up to five predictors of each outcome. The best model was chosen based on the combination of the highest adjusted R^2^, lowest Bayesian Information Criterion (BIC), lowest Akaike Information Criterion (AIC), and lowest Mallows’ Cp. Because each of these criteria may favour different models, and to address the issue of potential overfitting, we also considered the criterion of the lowest prediction error (CV) based on five-fold cross-validation (Lever et al., 2016). That is, we divided the data set into five subsets, whereby each subset (20%) served as test data and the remaining subsets (80%) as training data. The coefficients and related statistics for the chosen predictors of the best fits regression models for all outcome variables are summarised in Table 3. In the text we also reported adjusted R^2^, AIC, and CV as the most consistent indicators of the best model fits.

**Table 3.**
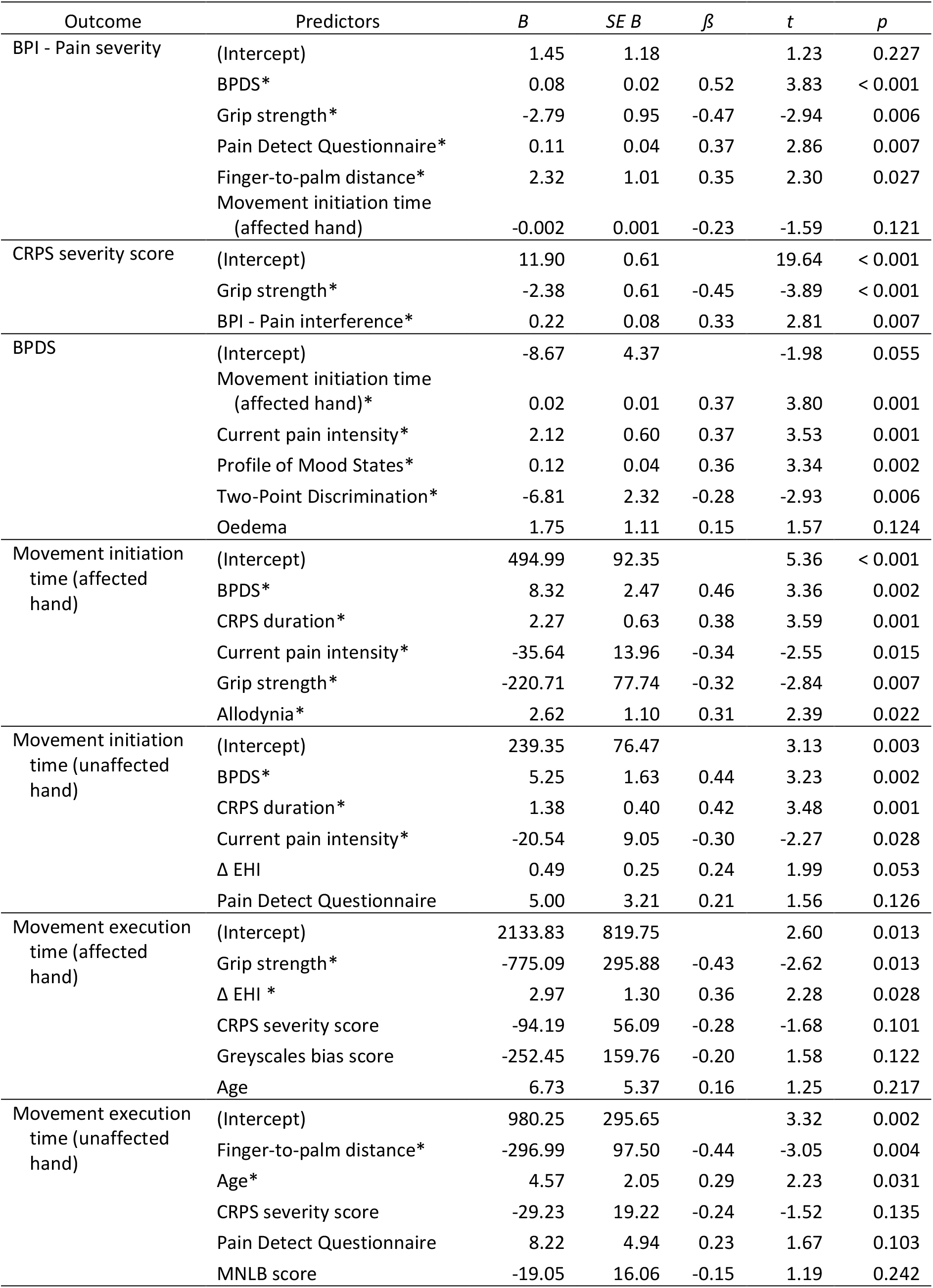
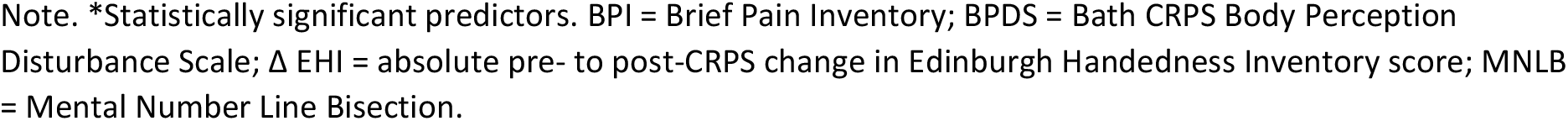
Model summaries for best subsets regression analyses

| Outcome | Predictors | *B* | *SE B* | $\beta$ | *t* | *p* |
| --- | --- | --- | --- | --- | --- | --- |
| BPI - Pain severity | (Intercept) | 1.45 | 1.18 |  | 1.23 | 0.227 |
|  | BPDS* | 0.08 | 0.02 | 0.52 | 3.83 | < 0.001 |
|  | Grip strength* | -2.79 | 0.95 | -0.47 | -2.94 | 0.006 |
|  | Pain Detect Questionnaire* | 0.11 | 0.04 | 0.37 | 2.86 | 0.007 |
|  | Finger-to-palm distance* | 2.32 | 1.01 | 0.35 | 2.30 | 0.027 |
|  | Movement initiation time (affected hand) | -0.002 | 0.001 | -0.23 | -1.59 | 0.121 |
| CRPS severity score | (Intercept) | 11.90 | 0.61 |  | 19.64 | < 0.001 |
|  | Grip strength* | -2.38 | 0.61 | -0.45 | -3.89 | < 0.001 |
|  | BPI - Pain interference* | 0.22 | 0.08 | 0.33 | 2.81 | 0.007 |
| BPDS | (Intercept) | -8.67 | 4.37 |  | -1.98 | 0.055 |
|  | Movement initiation time (affected hand)* | 0.02 | 0.01 | 0.37 | 3.80 | 0.001 |
|  | Current pain intensity* | 2.12 | 0.60 | 0.37 | 3.53 | 0.001 |
|  | Profile of Mood States* | 0.12 | 0.04 | 0.36 | 3.34 | 0.002 |
|  | Two-Point Discrimination* | -6.81 | 2.32 | -0.28 | -2.93 | 0.006 |
|  | Oedema | 1.75 | 1.11 | 0.15 | 1.57 | 0.124 |
| Movement initiation time (affected hand) | (Intercept) | 494.99 | 92.35 |  | 5.36 | < 0.001 |
|  | BPDS* | 8.32 | 2.47 | 0.46 | 3.36 | 0.002 |
|  | CRPS duration* | 2.27 | 0.63 | 0.38 | 3.59 | 0.001 |
|  | Current pain intensity* | -35.64 | 13.96 | -0.34 | -2.55 | 0.015 |
|  | Grip strength* | -220.71 | 77.74 | -0.32 | -2.84 | 0.007 |
|  | Allodynia* | 2.62 | 1.10 | 0.31 | 2.39 | 0.022 |
| Movement initiation time (unaffected hand) | (Intercept) | 239.35 | 76.47 |  | 3.13 | 0.003 |
|  | BPDS* | 5.25 | 1.63 | 0.44 | 3.23 | 0.002 |
|  | CRPS duration* | 1.38 | 0.40 | 0.42 | 3.48 | 0.001 |
|  | Current pain intensity* | -20.54 | 9.05 | -0.30 | -2.27 | 0.028 |
| | $\Delta$ EHI | 0.49 | 0.25 | 0.24 | 1.99 | 0.053 |
|  | Pain Detect Questionnaire | 5.00 | 3.21 | 0.21 | 1.56 | 0.126 |
| Movement execution time (affected hand) | (Intercept) | 2133.83 | 819.75 |  | 2.60 | 0.013 |
|  | Grip strength* | -775.09 | 295.88 | -0.43 | -2.62 | 0.013 |
| | $\Delta$ EHI * | 2.97 | 1.30 | 0.36 | 2.28 | 0.028 |
|  | CRPS severity score | -94.19 | 56.09 | -0.28 | -1.68 | 0.101 |
|  | Greyscales bias score | -252.45 | 159.76 | -0.20 | 1.58 | 0.122 |
|  | Age | 6.73 | 5.37 | 0.16 | 1.25 | 0.217 |
| Movement execution time (unaffected hand) | (Intercept) | 980.25 | 295.65 |  | 3.32 | 0.002 |
|  | Finger-to-palm distance* | -296.99 | 97.50 | -0.44 | -3.05 | 0.004 |
|  | Age* | 4.57 | 2.05 | 0.29 | 2.23 | 0.031 |
|  | CRPS severity score | -29.23 | 19.22 | -0.24 | -1.52 | 0.135 |
|  | Pain Detect Questionnaire | 8.22 | 4.94 | 0.23 | 1.67 | 0.103 |
|  | MNLB score | -19.05 | 16.06 | -0.15 | 1.19 | 0.242 |

Note. \*Statistically significant predictors. BPI = Brief Pain Inventory; BPDS = Bath CRPS Body Perception Disturbance Scale; $\Delta$ EHI = absolute pre- to post-CRPS change in Edinburgh Handedness Inventory score; MNLB = Mental Number Line Bisection.

##### 3.4.5.1. Predictors of pain and CRPS symptoms severity

The best fits regression models for the predictions of BPI pain severity and CRPS severity score are summarised in Table 3. Higher pain severity (as measured by BPI) was best predicted by more severe body perception disturbance, weaker grip strength in the affected hand, a greater neuropathic component of pain, greater range of movement in the affected hand, and faster movement initiation with the affected hand (non-significant predictor), *F*(5, 37) = 9.12, *p* < .001, *adj*. *R*^*2*^ = .49, AIC = 29.33, CV = 1.31. Larger CRPS severity scores were best predicted by weaker grip strength in the affected hand and higher pain interference, *F*(2, 51) = 20.59, *p* < .001, *adj. R*^*2*^ = .43, AIC = 26.30, CV = 1.22.

##### 3.4.5.2. Predictors of cognitive changes in CRPS

The best fits regression models for the predictions of BPDS and overall movement initiation and execution times with the affected and unaffected hands are summarised in Table 3. More severe body perception disturbance (higher BPDS score) was best predicted by slower movement initiation time with the affected hand, higher current pain intensity, higher mood disturbance score, more precise two-point discrimination on the affected limb, and greater swelling of the affected limb (non-significant predictor), *F*(5, 37) = 15.73, *p* < .001, *adj. R*^*2*^ = .64, AIC = 175.30, CV = 7.51. Slower initiation of movements with the affected hand was best predicted by more severe body perception disturbance, longer CRPS duration, lower current pain intensity, weaker grip strength in the affected hand, and more severe allodynia on the affected limb, *F*(5, 37) = 10.58, *p* < .001, *adj. R*^*2*^ = .53, AIC = 434.52, CV = 152.47. Movement initiation times with the unaffected hand shared some of the same predictors. Specifically, slower movement initiation was best predicted by more severe body perception disturbance, longer CRPS duration, lower current pain intensity, greater change in handedness after CRPS onset, and a greater neuropathic component of pain, although the latter two factors were not significant predictors, *F*(5, 44) = 6.59, *p* < .001, *adj. R*^*2*^ = .36, AIC = 475.81, CV = 129.60. Slower movement execution when using the affected hand was best predicted by weaker grip strength in the affected hand, greater change in handedness after CRPS onset, lower CRPS severity score, greater attention to the affected side of space on greyscales task, and older age, *F*(5, 39) = 4.81, *p* = .002, *adj. R*^*2*^ = .30, AIC = 559.36, CV = 492.89. However, only the first two factors were statistically significant predictors of movement execution time. For the unaffected hand, slower movement execution was best predicted by smaller range of movement in the affected hand, older age, lower CRPS severity score, greater neuropathic component of pain, and greater bias towards the affected side of the mental representation of space, *F*(5, 44) = 4.23, *p* = .003, *adj. R*^*2*^ = .25, AIC = 524.46, CV = 182.27. Only the first two factors significantly predicted movement execution time with the unaffected limb.

## 4. Discussion

We conducted a detailed examination of changes in spatial cognition in CRPS using sensitive experimental methods in a larger than previous research has used sample. Contrary to our hypotheses, our findings across measures of visuospatial attention and mental representation of space consistently showed no evidence of any spatial biases among people with CRPS compared to pain-free control participants, and there was very little evidence for directional motor deficits. We also found no support for any clinical relevance of changes in spatial cognition for the severity of pain and other symptoms of CRPS.

### 4.1. Visuospatial attention

Although previous studies have used TOJs to provide evidence for reduced tactile (Moseley et al., 2009, 2012; Reid et al., 2016) and visual (Bultitude et al., 2017; Filbrich et al., 2017) attention to the affected relative to unaffected side in people with CRPS, we found no such visuospatial attention bias on our TOJ task. One notable difference between these previous studies and ours is that most of them (Bultitude et al., 2017; Moseley et al., 2009; Reid et al., 2016) asked participants only to indicate which stimulus occurred *first*. This might mean that previous results were influenced by response bias, that is, a preference of one response over the other when the participant is uncertain about temporal order of the stimuli (Filbrich et al., 2016; Spence & Parise, 2010). Distorted perception of the CRPS-affected limb can involve hostile feelings, such as repulsion and hate (Lewis et al., 2007), which resemble misoplegia after brain injury (Bartolomeo et al., 2017). Thus, particularly when a verbal response is required (Bultitude et al., 2017), participants with CRPS might be reluctant to say “left” or “right”, depending on the side corresponding to their affected limb. Here we controlled for potential response bias by including a separate block of the TOJ in which participants were asked to indicate which stimulus occurred *second* (in addition to “which occurred first” block). The two previous studies that also controlled for response bias in a similar way, reported mixed findings regarding spatial biases on tactile TOJs (reduced attention to the affected side, Moseley et al., 2012, and normal performance, Filbrich et al., 2017). Thus the response bias might be an important factor in the performance of CRPS patients on the TOJ task. Consistent with our findings, Filbrich et al. (2017) found no apparent shift of visual attention when the participants’ hands were kept close to the trunk (outside of the visual field). However, when the stimuli appeared in immediate proximity of participants’ hands, they found a significant visuospatial bias even when controlling for response bias. This is in keeping with a proposal that spatial biases might only be present (or exacerbated) when body-relevant information is highly salient to the task (Reid et al., 2016). Thus, our results do not rule out the possibility that people with CRPS might still present with spatial biases in the tactile modality and / or related to other bodily information.

We also found that people with CRPS did not present with any biases on two additional tests of visual attention to and representation of near space. The Greyscales and Landmark tasks have not been previously tested in CRPS, but have sufficient sensitivity to detect visuospatial biases in brain-injured neglect patients (Mattingley et al., 2004), neurologically healthy individuals (Nicholls et al., 1999), and upper-limb amputees (Makin et al., 2010). Overall, consistently unbiased performance on the experimental tests of covert and overt attention to and representation of near visual space in this study suggests normal visuospatial cognition in CRPS. These findings agree with another study that also did not demonstrate any visuospatial biases in the speed of orienting saccades towards targets in either side of space (Filippopulos et al., 2015).

### 4.2. Mental representation of space

Representational neglect has not been extensively studied in CRPS, and not in combination with sensitive tests of visuospatial attention. One group study reported a shift of subjective midpoint of mental number line in the direction corresponding to the unaffected side (Sumitani et al., 2014), consistent with representational neglect after brain injury (Zorzi et al., 2002). However, a shift in the opposite direction was also reported in a single CRPS patient (Christophe, Delporte, et al., 2016; Jacquin-Courtois et al., 2017). While using the same MNLB task as Sumitani et al. (2014), we additionally presented number pairs not only in ascending, but also in descending order. Averaging the responses from these two conditions accounts for a potential tendency to report subjective midpoints as numbers closer to the starting point on mental number line, that is nearer the first number from a pair. Having controlled for these potential response biases, we did not find any systematic deviations from objective midline in participants with CRPS, nor any differences between the performance of participants with CRPS and controls.

### 4.3. Spatially-defined motor function

In general, there are four potential explanations of impaired motor function in CRPS. First, diagnostic criteria for CRPS include motor signs, such as weakness, decreased range of movement, or dystonia (Harden et al., 2010); thus physical pathology of the affected limb itself can result in impaired motor performance. Second, learned underutilization of the affected limb can develop through initial immobilization following a trauma, pain avoidance, and compensatory use of the unaffected limb (Punt et al., 2013). These learned behaviours can reinforce reduced use of the CRPS-affected limb and further deter its motor function. Third, motor “neglect-like” impairment can account for reduced or slower movements of the affected limb that cannot be attributed to any peripheral pathology, as well as movements performed in / towards the affected side of space, regardless of which limb is used (Laplane & Degos, 1983; Mattingley et al., 1992). Thus, motor function can be impaired in a spatially-defined manner consistent with neglect of the CRPS-affected limb and side of space. Fourth, central deficit of motor control can account for generalised / bilateral motor impairment that cannot be explained by peripheral pathology or deficits in spatial cognition. While the motor signs of CRPS and learned underutilization can only account for motor deficits specific to the CRPS-affected limb, the motor neglect and reorganization of central motor circuits additionally address spatially-defined and bilateral motor deficits found in CRPS, respectively. In the present study, we tried to dissect the motor neglect hypothesis from the alternative explanations of impaired motor function in CRPS.

Consistent with motor neglect of the affected side, people with CRPS previously reported having to focus their attention on the painful limb to move it (Galer et al., 1995; Galer & Jensen, 1999). Furthermore, their motor performance on speeded button pressing and circle drawing tasks was slower, more variable, and less accurate when they used the affected limb, and also when movements were performed in the affected side of space regardless of which hand they used (Reid et al., 2018). These findings suggest spatially-defined disruption of motor control that cannot be explained by physical pathology or learned underutilization of the affected limb (although the bilateral spatially-defined motor deficits did not replicate in another group study, Christophe, Chabanat, et al., 2016, and a case study, Christophe, Delporte, et al., 2016, which both used similar motor tasks). We tested for the first time if people with CRPS show *directional* motor neglect, that is, slowing of initiation (directional hypokinesia) or execution (directional bradykinesia) of movements directed towards the affected relative to unaffected side of space, regardless of which limb is used (Heilman, Bowers, & Watson, 1983; Mattingley, Bradshaw, & Phillips, 1992; Sapir et al., 2007).

People with CRPS in our study showed some evidence of slower execution of movements of the unaffected hand when they were directed towards the affected compared to unaffected side of space, consistent with hypothesised directional bradykinesia towards the affected side. This slowing cannot be attributed to perceptual neglect, as it occurred regardless of reaching to targets in the affected or unaffected side of space. Nor can it be attributed to physical pathology or learned underutilization of the CRPS-affected limb, as participants used the unaffected hand. On an individual level, some cases could be classified as showing consistent directional deficits towards the affected side with either hand, although the number was very few (< 10%). This is consistent with the finding that a relatively small proportion of brain-injured patients presented with directional hypokinesia towards the contralesional side when using their ipsilesional hand on the same task (17%, Sapir et al., 2007). However, in the present study a larger proportion of individuals with CRPS (16-19%) was actually classified as showing directional slowing towards the *un*affected side, but only when using the affected hand. Furthermore, on a group level, people with CRPS showed no other signs of directional hypokinesia or bradykinesia towards the affected or unaffected side for either hand that would differentiate them from pain-free controls. Most of the effects observed in both groups could be explained by general (non-directional) biomechanical constraints such as differences in movement pathways and crossing the body midline (see Supplementary Material). Therefore, when the differences between participants with CRPS and pain-free controls are considered as a whole, the results do not support the presence of directional motor deficits.

Although we did not find systematic evidence for directional motor deficits resembling motor neglect, our results demonstrate that people with CRPS had overall slowing of initiation and execution of the movements of both limbs as compared to pain-free controls. This suggests a central motor deficit that cannot be explained by peripheral pathology or learned underutilization of the CRPS-affected limb. Previous sensitive kinematic analyses also showed impairment on motor tasks performed with both hands (Schilder et al., 2012) or with the unaffected hand (Ribbers et al., 2002) in CRPS compared to pain-free individuals. Neuroimaging evidence suggests that these deficits could be related to altered central motor circuits, that is decreased inhibition of bilateral motor cortex (Juottonen et al., 2002; Schwenkreis et al., 2003), and its increased bilateral activation during movements of the affected hand relative to rest (Maihofner et al., 2007). Therefore, slowing of initiation and execution of movements with both limbs could be related to functional reorganization in cortical motor networks. However, we also cannot rule out that fatigue or analgesic medication could contribute to an overall decrease in psychomotor speed in individuals with CRPS, although existing evidence does not support the latter alternative (Kendall et al., 2010; Landrø et al., 2013).

### 4.4. Relationships between clinical signs, motor deficits, and neuropsychological changes in participants with CRPS

An additional aim of our study was to identify any relationships between clinical signs, motor deficits, and cognitive / psychological changes of our participants with CRPS. Recognizing that our analyses were exploratory, we offer only tentative explanations of the observed effects that should be tested in further research. Across different clinical and experimental measures, performance of CRPS participants on our battery of spatial cognition tests did not contribute to the prediction of the clinical outcomes. Therefore, these neuropsychological changes might not pertain to the clinical signs of CRPS, which calls into question the potential benefits of neurocognitive treatments that target deficits in spatial cognition (e.g. prism adaptation, Torta, Legrain, Rossetti, & Mouraux, 2016). In fact, on the whole, the key clinical measures of pain and CRPS severity were predicted by other clinical measures. Specifically, both pain and CRPS symptom severity were predicted by weaker grip strength, and pain was additionally predicted by reduced range of movement in the affected hand, highlighting the relevance of motor impairment. More severe sensory abnormalities consistent with features of neuropathic pain also predicted greater pain severity. Furthermore, we found a relationship between the severity of CRPS symptoms and the extent to which pain interfered with daily life, including work, social life, mobility, sleep, or mood (however, as pain interference was co-linear with pain severity, CRPS severity could be related to either).

From all the measures that could imply cortical reorganisation relevant to higher cognition, only self-reported body perception disturbance (BPDS scores) was related to pain severity and motor function. The BPDS measures subjective ownership of the affected limb; awareness of its position; attention to and valence of feelings towards the painful extremity; as well as perceived distortions of its size, shape, and / or weight (Lewis & McCabe, 2010). Higher pain intensity was previously linked to reporting greater distortions of body representation, both on the BPDS (Lewis & Schweinhardt, 2012) and neglect-like symptoms questionnaire (which measures partly overlapping construct of body ownership; Frettlöh et al., 2006; Wittayer et al., 2018). Distorted cognitive representation of the affected limb could reflect reorganization in the somatosensory cortical areas corresponding to that limb. People with weaker activation in the somatosensory cortex contralateral to the CRPS-affected hand (Pleger et al., 2006) and those with greater body perception disturbance (Lewis & Schweinhardt, 2012) had worse tactile discrimination abilities on the affected hand and higher levels of pain. Our analyses showed that, in addition to pain and tactile discrimination thresholds, greater body perception disturbance was also predicted by greater mood disturbance. This is in line with previously demonstrated relationships between psychological distress and scores on the neglect-like symptoms questionnaire (Michal et al., 2016; Wittayer et al., 2018).

Overall slowing of movements was the only outcome from our battery of spatial cognition and motor function tests that differentiated people with CRPS from healthy controls. We found that those with slower movement initiation with the affected and unaffected hands had more severely distorted body perception. This suggests that higher-order cognitive representations can contribute to motor function in CRPS. Body representation relies on combined proprioceptive, vestibular, somatosensory, and visual information that interact with the motor control system to guide actions (Head & Holmes, 1912). Higher scores on the neglect-like symptoms questionnaire (which, like the BPDS, also regards disownership of the affected limb; Frettlöh et al., 2006) were previously linked to greater motor impairment and disability in individuals with CRPS (Kolb et al., 2012). While these distortions in body perception primarily concern the affected limb, arm position sense (which relies on proprioception) has been found to be impaired bilaterally in CRPS (Brun et al., 2019; Lewis et al., 2010). Thus, deficits in proprioception in both limbs might slow down movement initiation due to uncertainty about their current positions. Slowing of movement initiation with both limbs was also predicted by longer CRPS duration, consistent with the idea that central mechanisms would have greater contribution to CRPS symptomatology in more chronic stages of the disease (Birklein & Schlereth, 2015; Bruehl & Chung, 2015; Veldman et al., 1993). We also found that people with more weakness in the affected hand and greater change in hand preference following CRPS onset (taken as an approximation of functional impact of CRPS) were slower to initiate and execute movements with the affected extremity. This is consistent with the “learned non-use” hypothesis (Punt et al., 2013): that ongoing underutilization of the CRPS-affected limb leads to atrophy, muscle weakness, and movement slowing, further exacerbating or maintaining motor deficits. Taken together, our results suggest that not only functional underutilization of the affected limb, but also bilateral central mechanisms of motor control and body perception, might contribute to the extent of motor impairment in CRPS.

Overall, our exploratory analyses do not support the conclusion that changes in spatial cognition are relevant for the manifestation and severity of CRPS symptoms. Instead, body representation and motor abilities appear to be important determinants of CRPS pain and symptom severity.

### 4.5. Strengths and limitations

Our results suggest that previously reported “neglect-like” changes in spatial cognition in CRPS might have been overstated. There are several advantages of the present study that strengthen our confidence in this conclusion. We systematically tested for any visuospatial and spatially-defined motor biases using a battery of sensitive tests in a group of people with CRPS that was two-to-five times larger than tested before on the TOJs or spatial motor tasks. One possible reason for the disparity between our results and those of previous studies is that there are individual differences in the extent to which cognitive function is affected in CRPS. Considering the high variability in the clinical presentation of CRPS, different trajectories of symptom development and strategies to deal with pain might lead to distinct patterns of cognitive changes (Marinus et al., 2011). In other words, in heterogeneous conditions such as CRPS, effects might arise in small sample studies that may not replicate, potentially due to the chance selection of more individuals who happen to present with a certain deficit. Consistent with this account of variability in past results, our participants with CRPS showed larger range of individual bias scores on the spatial tasks than the controls (Figure 2), although this could also be partly because of the smaller sample size for the control group. Participants with CRPS also presented with greater variability in the direction of any spatial biases than has been previously assumed (there is only one reported CRPS case of increased attention to the affected side of near space: Christophe, Delporte, et al., 2016; Jacquin-Courtois et al., 2017). Our sample was not large enough to define any subgroups of people with CRPS who might present with spatial biases more extreme than those found in pain-free participants. We were not able to identify any common characteristics of those people who did show larger biases. This is partly due to the small number of such cases, as well as within-participant variability (the performance of participants with CRPS only moderately correlated between different spatial tasks). One exception is that most of the extreme spatial biases appeared to be consistent with a leftward bias (exaggerated pseudoneglect), regardless of the CRPS-affected side (see Supplementary Figure 3). While this phenomenon was previously observed in CRPS (Reinersmann et al., 2012; Verfaille et al., 2019), in the present study it was not significant on a group level when compared to pain-free controls. Overall, since none of the measures of spatial cognition showed any systematic biases in CRPS, nor were they related to pain intensity or CRPS severity, their prevalence and clinical relevance are questionable.

Another strength of our study is that we controlled for potential response biases, which might have contributed to seemingly significant biases in previous TOJ studies. We also accounted for the fact that spatial attention might not normally be evenly distributed across space (see pseudoneglect, Jewell & McCourt, 2000) by obtaining comparative data from pain-free individuals. Follow-up Bayesian analyses showed anecdotal-to-moderate evidence of no differences between CRPS and pain-free participants on the visuospatial tasks (see also confidence intervals in Figure 2 illustrating no deviation from zero).

Nonetheless, although we aimed to create a diverse battery of tests of spatial cognition, there are two limitations that might have prevented us from detecting previously-reported spatial biases in our participants with CRPS. First, we were unable to include measures of tactile attention or egocentric reference frame, two measures upon which biased performance has been previously reported in CRPS (Moseley et al., 2009, 2012; Reid et al., 2016; Reinersmann et al., 2012; Sumitani, Shibata, et al., 2007; Uematsu et al., 2009). This is because we designed our protocol such that it only required transportable equipment and thus could be administered at patients’ homes and in different research centres, in order to obtain large and representative sample. Second, most of our tasks did not involve body-relevant information, although it has been proposed that this might be critical for the manifestation of spatial biases in CRPS (Reid et al., 2016). The exception is our motor task, which by definition involves the body, and which revealed very little evidence of any systematic spatial deficits. Considering the above mentioned limitations, we cannot rule out the possibility that our participants with CRPS might have presented with deficits in other domains of spatial cognition than those assessed in our study.

The third limitation to the extent to which our results can be compared to those of previous studies that reported changes in spatial cognition in CRPS is that the duration of CRPS in our sample was on average longer (except when compared to Bultitude et al., 2017; 4 years vs. <1-3 years, Filbrich et al., 2017; Moseley et al., 2012, 2009; Reid et al., 2018, 2016; Sumitani et al., 2014). However, several arguments suggest that changes in spatial cognition should not become *less* apparent over time: (a) there are clinical indications of greater contribution of central mechanisms to the manifestation of CRPS in its more chronic stages (Birklein & Schlereth, 2015; Bruehl & Chung, 2015; Veldman et al., 1993); (b) we found that longer CRPS duration predicted bilateral slowing of movement initiation, consistent with central changes in motor circuits; (c) there is evidence of positive correlations between CRPS duration and the extent of body perception distortion, body-related visuospatial bias, and spatially-defined motor bias (Lewis & Schweinhardt, 2012; Moseley, 2004; Reid et al., 2016, 2018); and (d) other studies (Bultitude et al., 2017; Filbrich et al., 2017; Frettlöh et al., 2006; Michal et al., 2016; Reid et al., 2016; Reinersmann et al., 2012) found no relationship between CRPS chronicity and any biases in spatial cognition, including our own findings from spatial tasks (Pearson’s *rs* = 0.06 to 0.27; see Supplementary Figure 2). Another factor that could limit the extent to which our findings are comparable to previous experimental studies on spatial cognition in CRPS is that pain intensity reported by our participants was on average greater (except when compared to Bultitude et al., 2017, and Sumitani et al., 2014; 5.8/10 vs. 4.3-4.8/10, Filbrich et al., 2017; Moseley et al., 2012, 2009; Reid et al., 2018). However, previous research reported either positive relationships between pain intensity and severity of “neglect-like” symptoms (Frettlöh et al., 2006; Reid et al., 2016; Wittayer et al., 2018), or found no relationships between these factors (Bultitude et al., 2017; Filbrich et al., 2017; Michal et al., 2016; Moseley et al., 2009; Reid et al., 2016), including our own results (Pearson’s *rs* = −0.12 to 0.20; see Supplementary Figure 2). Therefore, it is unlikely that the longer average disease duration or greater average pain intensity in our sample compared to previous research prevented us from detecting any impairments in spatial cognition.

## 5. Conclusions

Overall, the present findings suggest that unilateral upper-limb CRPS does not disrupt visual attention, mental representations, or motor function in a spatially-defined manner, and thus counter the analogy between CRPS and hemispatial neglect after brain injury. Although there were no behavioural indications of central changes in brain networks governing spatial cognition, bilateral slowing of movements implies impairment of central mechanisms of motor control. These appear to be related to some of the clinical features of CRPS rather than any spatial biases, although the extent of distorted cognitive representation of the affected limb seems to play a role in movement initiation speed and pain severity. These results support the promotion of treatments that aim to normalize body perception and improve motor function.

## Availability of data and materials

Anonymised individual-level data generated during the current study (https://osf.io/6nkxf/ DOI 10.17605/OSF.IO/6NKXF), digital study materials (PsychoPy experiment files and experimental stimuli; https://osf.io/7fk2v/ DOI 10.17605/OSF.IO/7FK2V) and analysis scripts (https://osf.io/bdnp6/ DOI 10.17605/OSF.IO/BDNP6) have been archived in a publicly available OSF repository.

## Acknowledgements

This research was funded by Regional Sympathetic Dystrophy Syndrome Association.

