## Supplementary Material for "Disputing space-based biases in unilateral complex regional pain syndrome"

### Comorbidities and treatments of participants with CRPS

A small proportion of participants reported CRPS symptoms in body parts other than the primarily affected upper limb - most commonly the ipsilateral lower limb (7% of total sample), with single instances of contralateral lower limb (not meeting the CRPS diagnostic criteria), and both lower limbs. A quarter of the CRPS sample reported other, non-CRPS pain. In most cases it was fibromyalgia (13% of total sample), but there were also single instances of joint hypermobility; shoulder, hip, and back pain; migraine; hernia; peripheral neuropathy in the ipsilateral lower limb; and contralateral upper limb pain. The most common comorbidities other than pain included depression (35% of total sample), anxiety (20%), asthma (13%), hypertension (9%), polycystic ovaries (7%), diabetes, irritable bowel syndrome, and arthritis (6%). There were also single cases of hypothyroidism, tachycardia, endometriosis, psoriasis, contralateral carpal tunnel syndrome, epilepsy, anaemia, incontinence, Fowler's syndrome, and Crohn's disease. Ongoing treatments and medications for CRPS at the time of the study involved opioids (56%), anti-depressants (48%), anticonvulsants (46%), paracetamol (44%), physiotherapy and/or occupational therapy (39%), nonsteroidal anti-inflammatory drugs (33%), local anaesthetics (15%), other medication (6%), spinal cord stimulation (4%), and transcutaneous electrical nerve stimulation (4%).

### Spatially-defined motor function - results

The results of the linear mixed models analyses on the spatially-defined motor function task data are reported in Supplementary Table 1. In the main text, we reported the main effects of Group on movement initiation times with either hand, indicating that participants with Complex Regional Syndrome (CRPS) were overall slower compared to controls.

In addition to the above effects reported in the manuscript, there was also a main effect of Starting Position on initiation times with the affected limb suggesting that, regardless of Group or Visual Field (VF), movement initiation was slower from the affected (*Mdn* = 477.32, BCa 95% CI [455.98, 532.44]) than unaffected side of space (*Mdn* = 458.88, BCa 95% CI [435.17, 505.13]). This pattern is consistent with directional hypokinesia towards the unaffected side (i.e. slowing of movements directed toward the unaffected, rather than affected side of space).

Also reported in the main text is that main effects of Group were also found for movement execution times with either hand, with slowing among CRPS patients relative to controls. There was also an interaction between Group and hand Starting Position for the unaffected limb, suggesting slower execution of movements from the unaffected than affected Starting Position among CRPS patients compared to controls.

In addition to the above results reported in the main text, we found the following effects on execution times that did not involve Group. A main effect of VF suggested that, independent of Group or Starting Position, execution of movements with the affected limb was slower towards the targets in the unaffected (*Mdn* = 891.65, BCa 95% CI [795.69, 939.31]) compared to the affected (*Mdn* = 826.69, BCa 95% CI [770.99, 924.28]) side of space. This suggests that movements to the targets in the side of space ipsilateral to the affected limb were faster than to the targets in the contralateral side of space.

Main effects of Starting Position also indicated overall slower movement execution from the affected (*Mdn* = 937.65, BCa 95% CI [810.78, 980.04]) than central (*Mdn* 799.22, BCa 95% CI [743.69, 880.67]) position with the affected limb, and from the unaffected (*Mdn* = 786.88, BCa 95% CI [739.55, 847.88]) than affected (*Mdn* = 750.09, BCa 95% CI [707.51, 803.14]) position with the unaffected limb, regardless of Group or VF. These suggest that movements from starting positions ipsilateral to the hand used were slower than from contralateral or central positions.

Furthermore, there were interactions between Starting Position and VF on execution times (Supplementary Figure 1). When executing movements with the affected or unaffected hand, participants were slower in reaching targets in the unaffected VF from the affected Starting Position (affected hand *Mdn* = 943.60, BCa 95% CI [847.43, 1047.72]; unaffected hand *Mdn* = 789.42, BCa 95% CI [702.34, 824.93]) than affected VF (affected hand *Mdn* = 887.55, BCa 95% CI [790.22, 926.11]; unaffected hand *Mdn* = 726.93, BCa 95% CI [681.03, 774.97]; Supplementary Figure 1a) . Conversely participants were slower in reaching targets in the affected VF from the unaffected Starting Position (affected hand *Mdn* = 857.01, BCa 95% CI [781.54, 931.30]; unaffected hand *Mdn* = 821.87, BCa 95% CI [776.09, 860.07]) than unaffected VF (affected hand *Mdn* = 814.83, BCa 95% CI [754.59, 905.28]; unaffected hand *Mdn* = 759.12, BCa 95% CI [705.55, 802.46]; Supplementary Figure 1b). Movement execution with the affected hand towards the targets in the unaffected VF was significantly slower from the affected than unaffected Starting Position (Supplementary Figure 1c), whereas when the unaffected hand was used, movement execution to the targets in the affected VF was significantly slower from the unaffected than affected Starting Position (Supplementary Figure 1d). These interactions suggest that regardless of Group, participants were slower to execute the longer movements that crossed the body midline with either hand than to execute the shorter movements that were made within the same side of the body midline. Additionally, participants were slower to execute movements from the central Starting Position with the unaffected hand towards the affected VF (*Mdn* = 717.05, BCa 95% CI [679.23, 802.12]) than unaffected VF (*Mdn* = 690.13, BCa 95% CI [637.05, 754.59]), regardless of Group (Supplementary Figure 1e). This effect would be consistent with directional bradykinesia towards the affected side. Alternatively, it could reflect slowing of movement to the side of space contralateral to the limb used. However, execution of the unaffected hand movements towards the unaffected VF (that is, side of space ipsilateral to the limb used) was also slower from the affected than central Starting Position (Supplementary Figure 1f). There were no further significant main effects or interactions for movement execution times.

Overall, the results provide some evidence for directional bradykinesia towards the affected side when using the unaffected limb, as well as directional hypokinesia and bradykinesia towards the unaffected side when using the affected limb. However, these patterns were not specific to CRPS, but largely consistent across both groups. Most of the significant effects for movement execution time as a function of hand starting position and target location can be explained by the length of the required movement pathway and whether it crossed the body midline. In summary, we observed little systematic evidence of directional motor deficits consistent with motor neglect in participants with CRPS.

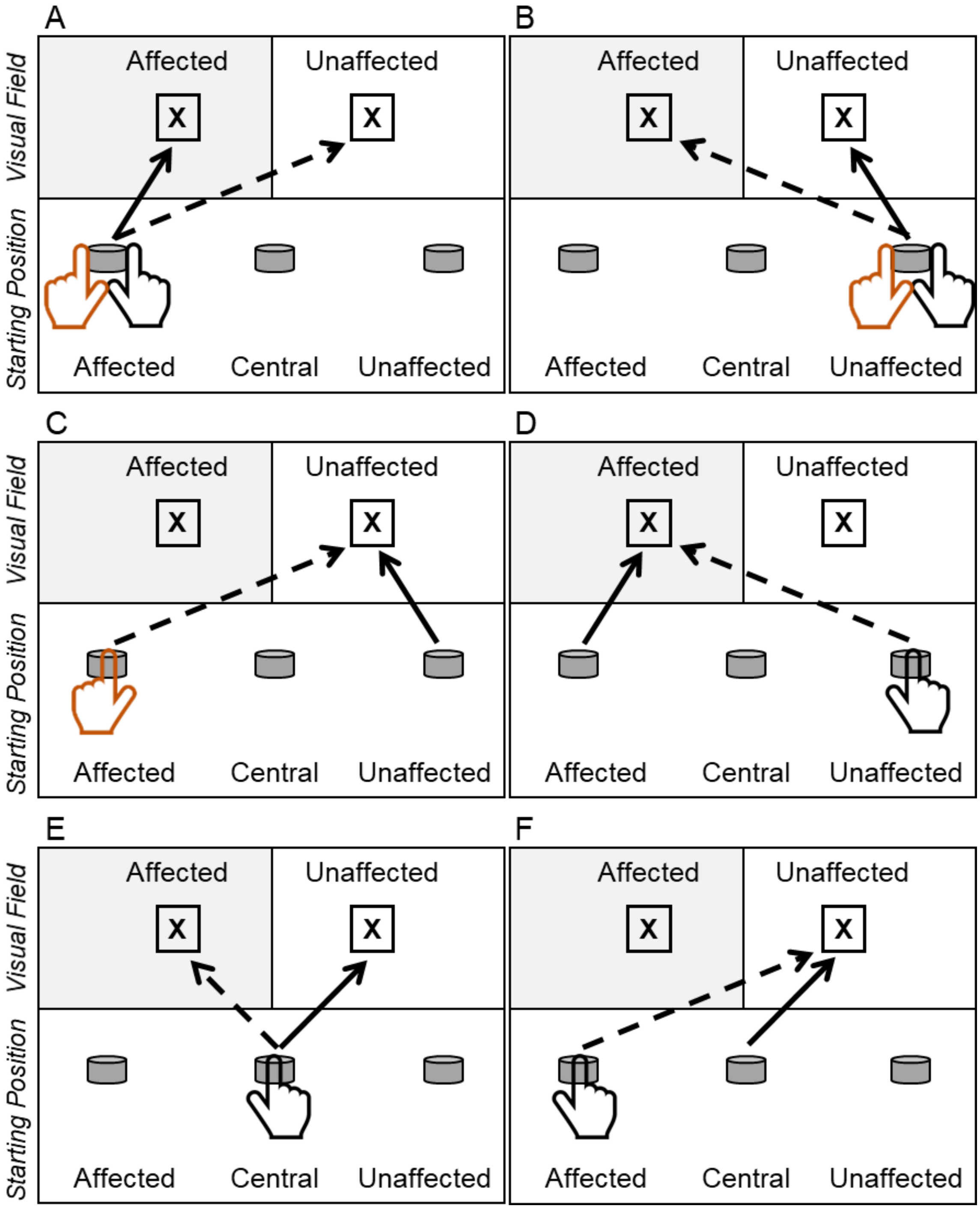


*Supplementary Figure 1.* Diagrams illustrate the results of significant post-hoc contrasts following Starting Position x Visual Field interactions on movement execution times with the affected (orange) and unaffected (black) hand. Arrows indicate the direction of movement. Dashed lines represent slower movement execution as compared to solid lines. “X” represents the target, and the grey cylinders represent the possible button locations / hand Starting Positions. Regardless of Group, participants were slower to execute movements (A) towards the unaffected VF than affected VF from the affected Starting Position, with either hand; (B) towards the unaffected VF than affected VF from the unaffected Starting Position, with either hand; (C) towards the unaffected VF from the affected than unaffected Starting Position, with the affected hand; (D) towards the affected VF from the unaffected than affected Starting Position, with the unaffected hand; (E) towards the affected VF than unaffected VF from the central Starting Position, with the unaffected hand; (F) towards the unaffected VF from the affected than central Starting Position, with the unaffected hand.

Supplementary Table 1

*The results of bootstrapped (n = 1000) linear mixed modelling of main effects and interactions on movement initiation / execution times for each hand in the test of spatially-defined motor function*

| Model term | Movement initiation time Affected hand | | | Movement initiation time Unaffected hand | | | Movement execution time Affected hand | | | Movement execution time Unaffected hand | | |
|---|---|---|---|---|---|---|---|---|---|---|---|---|
| | Regression coefficient | Lower 95% CI | Upper 95% CI | Regression coefficient | Lower 95% CI | Upper 95% CI | Regression coefficient | Lower 95% CI | Upper 95% CI | Regression coefficient | Lower 95% CI | Upper 95% CI |
| Intercept | 0.433* | 0.425 | 0.440 | 0.421* | 0.413 | 0.429 | 0.693* | 0.674 | 0.713 | 0.672* | 0.641 | 0.702 |
| Group (Con vs CRPS) | 0.191* | 0.159 | 0.222 | 0.073* | 0.049 | 0.099 | 0.444* | 0.357 | 0.523 | 0.112* | 0.077 | 0.147 |
| Starting Position (AP vs CP) | -0.009 | -0.021 | 0.003 | 0.006 | -0.005 | 0.019 | -0.083* | -0.112 | -0.057 | 0.006 | -0.040 | 0.054 |
| Starting Position (AP vs UP) | -0.025* | -0.036 | -0.012 | 0.006 | -0.008 | 0.020 | -0.006 | -0.044 | 0.029 | 0.066* | 0.015 | 0.118 |
| Visual Field (AVF vs UVF) | -0.005 | -0.016 | 0.007 | -0.007 | -0.019 | 0.004 | 0.050* | 0.021 | 0.078 | 0.021 | -0.021 | 0.061 |
| Group (Con vs CRPS) x Starting Position (AP vs CP) | -0.008 | -0.049 | 0.038 | -0.020 | -0.055 | 0.008 | -0.010 | -0.109 | 0.085 | 0.007 | -0.044 | 0.061 |
| Group (Con vs CRPS) x Starting Position (AP vs UP) | 0.010 | -0.035 | 0.055 | -0.008 | -0.043 | 0.023 | 0.047 | -0.064 | 0.164 | 0.063* | 0.002 | 0.120 |
| Group (Con vs CRPS) x Visual Field (AVF vs UVF) | 0.023 | -0.021 | 0.068 | -0.010 | -0.041 | 0.018 | 0.110 | -0.003 | 0.222 | 0.020 | -0.029 | 0.068 |
| Starting Position (AP vs CP) x Visual Field (AVF vs UVF) | 0.008 | -0.010 | 0.024 | 0.004 | -0.013 | 0.022 | -0.023 | -0.057 | 0.012 | -0.066* | -0.126 | < -0.001 |
| Starting Position (AP vs UP) x Visual Field (AVF vs UVF) | 0.009 | -0.008 | 0.026 | 0.006 | -0.012 | 0.025 | -0.085* | -0.129 | -0.040 | -0.079* | -0.143 | -0.014 |
| Group (Con vs CRPS) x Starting Position (AP vs CP) x Visual Field (AVF vs UVF) | -0.005 | -0.062 | 0.052 | -0.004 | -0.041 | 0.037 | 0.060 | -0.123 | 0.284 | -0.022 | -0.099 | 0.052 |
| Group (Con vs CRPS) x Starting Position (AP vs UP) x Visual Field (AVF vs UVF) | -0.005 | -0.077 | 0.068 | 0.011 | -0.025 | 0.050 | -0.105 | -0.264 | 0.051 | -0.036 | -0.109 | 0.042 |

*Note.* *Significant effect (95% CI around the regression coefficient estimate does not include zero). The levels of each model term are indicated in brackets, with the reference term listed first. Group factor had two levels (Con = control participants, CRPS = participants with Complex Regional Pain Syndrome) and Visual Field factor also had two levels (AVF = affected, UVF = unaffected). The hand Starting Position factor had three levels (CP = central, AP = affected, UP = unaffected), necessitating the inclusion of two terms in the model for each main effect and interaction involving this Factor.

| | CSS | Current pain | BPI severity | BPI interference | Age | Duration | PDQ | Tampa | BPDS | POMS | EHI (pre-CRPS) | Abs. ΔEHI | Temp. difference | Abs. temp. difference | Abs. temp. difference | Grip strength | ΔFTP | MDT | MPT | Allodynia | Two-point discrim. | PSS | PSE | Greyscales | MNLB | MIT AH | MIT UH | MET AH |
|---|---|---|---|---|---|---|---|---|---|---|---|---|---|---|---|---|---|---|---|---|---|---|---|---|---|---|---|---|
| Current pain | **0.30** | | | | | | | | | | | | | | | | | | | | | | | | | | | |
| BPI severity | **0.43** | **0.90** | | | | | | | | | | | | | | | | | | | | | | | | | | |
| BPI interference | **0.53** | **0.57** | 0.73 | | | | | | | | | | | | | | | | | | | | | | | | | |
| Age | -0.19 | -0.12 | -0.17 | -0.11 | | | | | | | | | | | | | | | | | | | | | | | | |
| Duration | -0.12 | 0.11 | 0.07 | 0.08 | **0.30** | | | | | | | | | | | | | | | | | | | | | | | |
| PDQ | **0.43** | **0.38** | **0.45** | **0.50** | -0.07 | -0.13 | | | | | | | | | | | | | | | | | | | | | | |
| Tampa | 0.22 | 0.22 | **0.30** | **0.37** | -0.05 | 0.09 | 0.22 | | | | | | | | | | | | | | | | | | | | | |
| BPDS | **0.34** | **0.49** | **0.52** | **0.46** | -0.07 | -0.07 | **0.47** | **0.39** | | | | | | | | | | | | | | | | | | | | |
| POMS | 0.15 | 0.24 | **0.34** | **0.46** | -0.16 | -0.09 | **0.44** | **0.51** | **0.59** | | | | | | | | | | | | | | | | | | | |
| EHI (pre-CRPS) | 0.08 | -0.09 | -0.08 | 0.13 | 0.24 | 0.06 | 0.15 | 0.11 | 0.19 | 0.11 | | | | | | | | | | | | | | | | | | |
| Abs. ΔEHI | **0.46** | 0.04 | 0.20 | 0.24 | -0.12 | -0.21 | 0.19 | 0.21 | 0.16 | 0.04 | 0.08 | | | | | | | | | | | | | | | | | |
| Temp. difference | 0.02 | 0.22 | 0.24 | 0.23 | 0.07 | **0.29** | 0.14 | **0.40** | 0.20 | 0.25 | 0.16 | 0.05 | | | | | | | | | | | | | | | | |
| Abs. temp. difference | 0.14 | -0.14 | -0.10 | -0.09 | -0.02 | -0.16 | -0.12 | **-0.39** | -0.16 | **-0.30** | -0.17 | 0.16 | **-0.84** | | | | | | | | | | | | | | | |
| Oedema | 0.09 | **-0.32** | -0.25 | -0.02 | 0.06 | -0.22 | -0.05 | -0.03 | -0.04 | -0.09 | 0.22 | **0.48** | 0.03 | 0.05 | | | | | | | | | | | | | | |
| Grip strength | **-0.60** | **-0.33** | **-0.44** | **-0.45** | 0.08 | 0.07 | -0.24 | -0.10 | **-0.33** | -0.19 | -0.11 | **-0.45** | -0.03 | -0.17 | 0.17 | | | | | | | | | | | | | |
| ΔFTP | **-0.47** | -0.14 | -0.25 | -0.24 | -0.09 | 0.16 | -0.20 | 0.01 | -0.21 | -0.03 | -0.05 | **-0.41** | -0.07 | -0.16 | 0.01 | **0.68** | | | | | | | | | | | | |
| MDT | -0.21 | -0.03 | -0.08 | 0.01 | 0.12 | 0.13 | -0.20 | -0.01 | -0.13 | -0.02 | 0.09 | 0.18 | **0.30** | -0.24 | 0.19 | 0.10 | -0.03 | | | | | | | | | | | |
| MPT | **0.29** | 0.18 | 0.19 | -0.02 | -0.11 | 0.02 | 0.22 | 0.16 | 0.23 | -0.13 | -0.06 | 0.00 | -0.10 | 0.11 | 0.00 | 0.00 | 0.04 | **-0.27** | | | | | | | | | | |
| Allodynia | **0.32** | **0.45** | **0.51** | **0.36** | -0.09 | -0.08 | **0.52** | 0.14 | **0.50** | **0.32** | 0.11 | 0.16 | -0.03 | 0.04 | -0.10 | **-0.38** | **-0.29** | **-0.38** | **0.28** | | | | | | | | | |
| Two-point discrim. | -0.09 | -0.02 | -0.08 | 0.07 | 0.01 | 0.04 | -0.02 | 0.07 | -0.04 | 0.17 | -0.20 | -0.08 | 0.04 | -0.11 | -0.14 | -0.06 | 0.03 | 0.05 | -0.17 | 0.09 | | | | | | | | |
| PSS | 0.18 | 0.07 | 0.16 | **0.27** | 0.01 | 0.12 | 0.20 | **0.30** | 0.20 | 0.13 | 0.06 | 0.01 | 0.20 | -0.21 | -0.02 | -0.08 | 0.04 | -0.18 | 0.20 | -0.08 | -0.17 | | | | | | | |
| PSE | 0.07 | 0.18 | 0.20 | **0.29** | 0.10 | **0.27** | 0.08 | 0.05 | 0.10 | 0.08 | 0.00 | -0.22 | 0.22 | 0.01 | **-0.30** | -0.13 | -0.22 | 0.15 | -0.16 | -0.14 | 0.00 | 0.10 | | | | | | |
| Greyscales | -0.11 | 0.05 | -0.12 | -0.15 | 0.02 | 0.06 | -0.07 | -0.04 | -0.01 | -0.11 | **-0.33** | **-0.27** | -0.07 | 0.13 | **-0.28** | -0.04 | -0.04 | -0.06 | 0.06 | -0.19 | 0.06 | -0.01 | **0.40** | | | | | |
| MNLB | -0.07 | 0.20 | 0.15 | -0.04 | 0.06 | 0.06 | 0.06 | -0.10 | -0.02 | 0.00 | **-0.27** | **-0.54** | -0.07 | 0.10 | **-0.49** | 0.04 | 0.00 | -0.22 | 0.07 | -0.01 | -0.15 | -0.09 | **0.38** | **0.37** | | | | |
| MIT AH | **0.31** | 0.17 | **0.32** | 0.28 | 0.21 | **0.32** | **0.41** | 0.19 | **0.52** | **0.33** | 0.23 | 0.27 | 0.23 | -0.16 | -0.04 | **-0.44** | -0.25 | -0.26 | 0.10 | **0.45** | 0.09 | 0.26 | 0.12 | -0.22 | -0.23 | | | |
| MIT UH | 0.21 | 0.06 | 0.15 | 0.12 | **0.32** | **0.28** | **0.32** | 0.20 | **0.45** | 0.14 | 0.24 | 0.26 | 0.20 | -0.11 | 0.21 | -0.21 | -0.20 | -0.07 | 0.18 | 0.21 | -0.26 | **0.35** | 0.12 | 0.02 | -0.24 | **0.71** | | |
| MET AH | 0.16 | 0.02 | 0.21 | 0.20 | 0.14 | 0.01 | 0.09 | 0.12 | 0.12 | 0.26 | -0.04 | **0.45** | 0.13 | -0.02 | 0.02 | **-0.45** | **-0.33** | -0.01 | -0.29 | 0.15 | 0.29 | 0.07 | -0.07 | -0.26 | -0.12 | **0.54** | **0.21** | |
| MET UH | 0.05 | 0.17 | 0.21 | -0.02 | **0.32** | 0.00 | 0.25 | 0.15 | 0.05 | -0.05 | 0.06 | **0.33** | 0.18 | -0.15 | -0.01 | -0.18 | **-0.39** | 0.05 | -0.07 | 0.08 | 0.00 | 0.00 | 0.07 | -0.11 | -0.07 | **0.42** | **0.45** | **0.38** |

| Pearson correlation |
|---|
| 1.00 |
| 0.90 |
| 0.80 |
| 0.70 |
| 0.60 |
| 0.50 |
| 0.40 |
| 0.30 |
| 0.20 |
| 0.10 |
| 0.00 |
| -0.10 |
| -0.20 |
| -0.30 |
| -0.40 |
| -0.50 |
| -0.60 |
| -0.70 |
| -0.80 |
| -0.90 |
| -1.00 |

*Supplementary Figure 2.* Correlation matrix illustrating relationships between participant characteristics; self-reported pain, fear of movement, body perception, and mood; sensory, motor, and autonomic function; and experimental measures of visuospatial attention, mental representation of space, and movement speed. The numbers represent the Pearson's correlation coefficient. The strength and direction of the correlations are colour-coded according to the legend on the right-hand side. Numbers in bold represent correlations significant at the level of $p < .05$. CSS = CRPS symptom severity score; BPI = Brief Pain Inventory; PDQ = Pain Detect Questionnaire; Tampa = Tampa Scale for Kinesiophobia; BPDS = Bath CRPS Body Perception Disturbance Scale; POMS = Profile of Mood States; EHI = Edinburgh Handedness Inventory; Abs. ΔEHI = absolute change in handedness index from before CRPS onset to current handedness; ΔFTP = Finger-To-Palm distance; MDT = Mechanical Detection Threshold; MPT = Mechanical Pain Threshold; PSS = Point of Subjective Simultaneity; PSE = Point of Subjective Equality; MNLB = Mental Number Line Bisection; MIT AH = movement initiation time with the affected hand; MIT UH = movement initiation time with the unaffected hand; MET AH = movement execution time with the affected hand; MET UH = movement execution time with the unaffected hand.

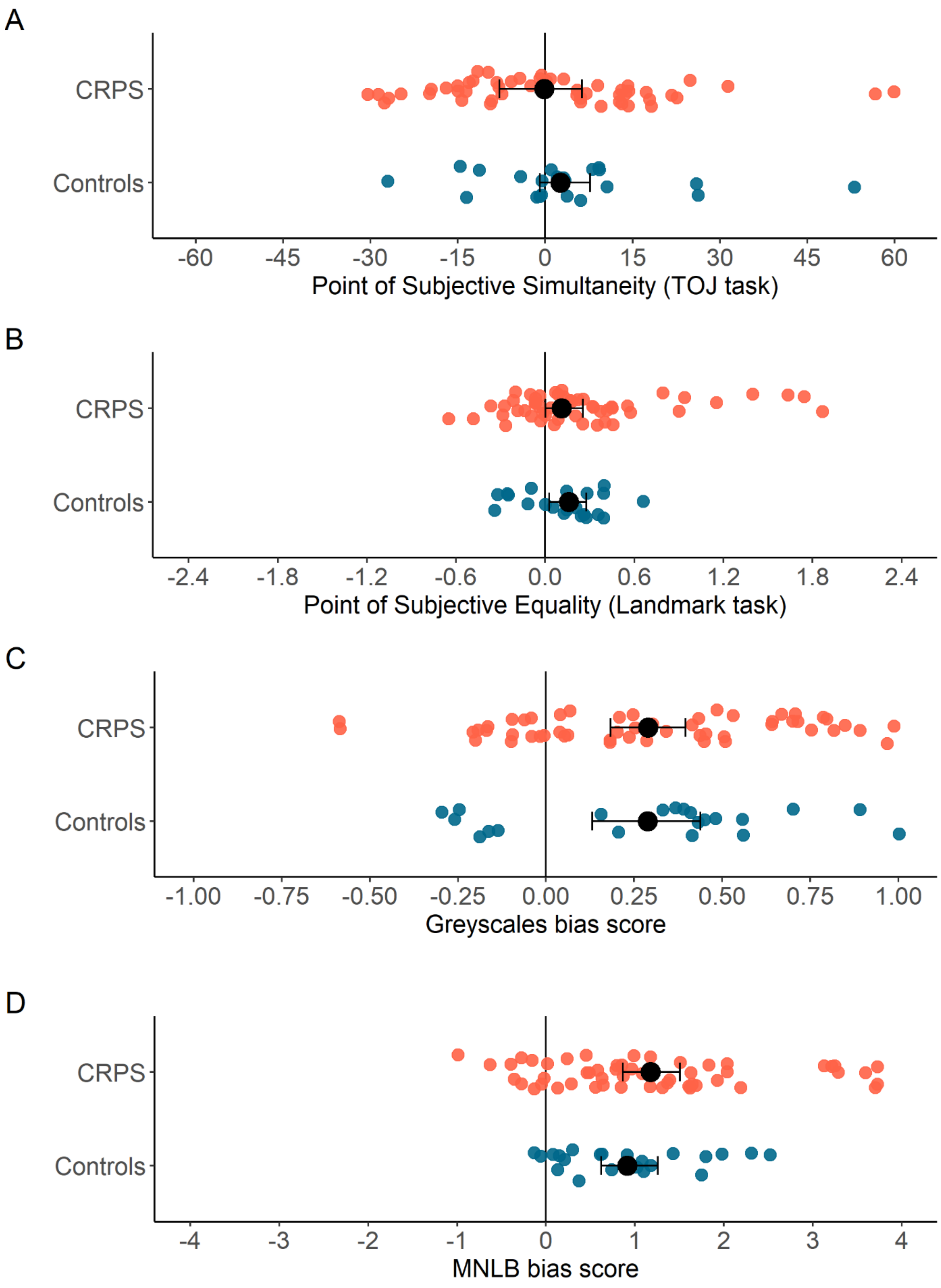


*Supplementary Figure 3.* Results of the experimental tests of visuospatial attention (A-C) and the mental representation of space (D) based on the data coded according to left/right reference (see Figure 2 in the main article for the data coded according to affected/unaffected reference). Smaller circles represent individual data from participants with CRPS (orange) and pain-free control participants (blue). Larger black circles represent the group median (A, B) and mean (C, D) scores with bootstrapped 95% confidence intervals (error bars). (A) The Point of Subjective Simultaneity on the Temporal Order Judgement task indicates by how many milliseconds the light on the left side should precede (negative values) or follow (positive values) the light on the right side for the two lights to be perceived as simultaneous. (B) The Point of Subjective Equality on the Landmark task indicates by how many degrees of visual angle the pair of landmarks should be offset from being truly equidistant to central fixation towards the left side (negative values) or towards the right side (positive values) for the two landmarks to be perceived as equidistant. (C) The bias score on the Greyscales task indicates to what extent the participants were basing their darkness judgements on the side right side of the stimuli (negative values) or on the left side of the stimuli (positive values). (D) The bias score on the Mental Number Line Bisection task indicates to what extent participants' subjective midpoint of the mental number line was shifted towards the higher numbers (corresponding to the right side; negative values) or towards the smaller numbers (corresponding to the left side; positive values). Negative scores for each of the measures depicted in this figure would indicate reduced attention to or (mental) representation of the left side of space
